# Patterned integrin–laminin adhesion coordinates epithelial collective cell migration

**DOI:** 10.1101/2025.09.12.675859

**Authors:** Anna Mertens, Nicola Moratscheck, Petra A Klemmt, Chaitanya Dingare, Marion Basoglu, Virginie Lecaudey

**Affiliations:** Department of Developmental Biology of Vertebrates, Institute of Cell Biology and Neuroscience, Goethe Universität Frankfurt, Frankfurt am Main, Germany; Developmental Biology, Institute for Biology I, Faculty of Biology, Albert-Ludwigs-Universität Freiburg, Freiburg im Breisgau, Germany; Electron microscopy facility, FB15, Goethe Universität Frankfurt, Frankfurt am Main, Germany; Klinik für Neurologie und Neurophysiologie, Universitätsklinikum Freiburg, Freiburg im Breisgau, Germany; Referat Biologische Sicherheit, Goethe Universität Frankfurt, Frankfurt am Main, Germany; Laboratory of Comparative Developmental Dynamics, Department of Genetics, University of Cambridge, Cambridge, UK

## Abstract

Collective cell migration is essential for development, tissue homeostasis, and disease progression. Although integrin-mediated adhesion is well characterized in single migrating cells *in vitro*, how integrins coordinate collective epithelial migration *in vivo* is less clear. Using the zebrafish posterior lateral line primordium (pLLP), we show that integrins α3 and α6 are differentially expressed in the trailing and leading regions of the pLLP, respectively, and that their combined loss induces excessive protrusive activity and slows migration. We further identify laminin α5 (Lama5) as a key basement membrane (BM) component underlying migrating pLLP cells. While loss of laminin α5 alone compromises BM integrity without impairing migration, simultaneous depletion of *lama5* and *itga6b* markedly decreases velocity, and ultimately blocks migration. These findings reveal a robust, redundant adhesion machinery that ensures persistent collective migration *in vivo* and highlight fundamental principles of epithelial dynamics relevant to development and disease.

## Introduction

The migration of cells and tissues is fundamental to many stages of life, from embryogenesis, tissue regeneration and immune surveillance to cancer progression ^1–3^. To generate migratory forces, cells push or pull on their surroundings. Originating from actomyosin contractions, intracellular forces need to be transmitted to the extracellular matrix (ECM) via focal adhesions (FA) ^4,5^. The classic model of cell migration is centered around cycles of FA assembly (adhesion), maturation (tension build-up) and release (tension release) ^6–8^. The core FA protein integrins are heterodimeric α/β transmembrane receptors that bind ECM ligands extracellularly and connect to the cytoskeleton via adaptor proteins. In mammals, 18 α- and 8 β-subunits assemble into 24 distinct integrins with distinct ligand preferences, including laminins, collagens and RGD-containing proteins ^6^. Tissue- and context-specific expression of integrins influences the balance of actomyosin and adhesion forces ^9^. This allows cells to navigate complex environments and adapt to varying mechanical and chemical properties, as is the case in tumor invasion and development ^10^. The result is a spectrum of migration modes – as single cells or collectives, highly-adhesive mesenchymal or low-adhesive amoeboid, and anything in between ^5,9^.

Integrin-ECM adhesion is especially critical during development ^6^. Specialized sheet-like ECMs, termed basement membranes (BMs) provide cells with a physical barrier along which to orient, migrate, differentiate and signal ^11–13^. Heterotrimeric α/β/γ laminins are one of the core components of BMs, initiating BM assembly ^14^ and directly interacting with integrin receptors ^15^. Laminins-111 and -511 are the first to be expressed and essential for mouse embryogenesis ^16–18^. Accordingly, loss of laminin γ1-chain (Lamc1) leads to a complete lack of BMs and inability to gastrulate ^19^. Mutations in Lama5 reveal roles for laminin α5 in limb formation, neural tube closure, kidney and lung morphogenesis, and hair cell development ^18,20^. LAMA5 also contributes to tumor progression ^21–23^. Likewise, loss of integrin-laminin adhesion underlies human diseases ranging from congenital muscular dystrophy (laminin α2-integrin α7β1 adhesion defects) ^24^ to kidney dysfunction and skin blistering due to malformations of the epidermal BM (integrin α3β1 mutations) ^25^.

Despite extensive work in two-dimensional (2D) *in vitro* systems ^26–28^, our understanding of how integrin-laminin adhesions function in the physical and biochemical complexity of three-dimensional (3D), *in vivo* environments remains incomplete ^9,29^. As a result, fundamental questions remain about how cell–ECM adhesion is organized and regulated *in vivo*, how integrin-mediated adhesion complexes respond to tissue-scale mechanical cues ^30^, and how these interactions contribute to coordinated collective migration events ^31^.

The zebrafish posterior lateral line primordium (pLLP) provides a powerful model to address these questions. This cohesive cluster of cells migrates along the trunk while simultaneously undergoing proliferation, differentiation and neuromast deposition, combining features of collective migration, epithelial remodeling and morphogenesis in a 3D *in vivo* environment ^32,33^. The pLLP exhibits a polarized organization with a pseudo-mesenchymal leading edge and an epithelial trailing region, where cells undergo apical constriction to form neuromast precursors ^34,35^. As the pLLP migrates, neuromasts are deposited sequentially and remain connected through a chain of interneuromast cells (INCs). Recently, Yamaguchi and colleagues demonstrated that a laminin γ1-containing BM, the FA proteins integrin β1 (Itgb1a/Itgb1b) and talin are required for efficient migration of the pLLP ^36^. Yet, the broader molecular machinery that mediates adhesion between the pLLP and its migratory substrate remains poorly defined.

Here, we dissect the roles of integrins and laminins during pLLP migration. We identify the epithelial integrins α3a (Itga3a), α3b (Itga3b) and α6b (Itga6b) as components of a front-rear polarized adhesion machinery, which function hierarchically and partially redundantly. We further show that laminin α5 is an essential component of the BM underlying migrating pLLP cells. While loss of *lama5* disrupts the integrity of the BM without significantly affecting migration, combined depletion of *lama5* and *itga6b* strongly impacts pLLP velocity and ultimately blocks its progression. Together, our findings uncover redundant integrin–laminin adhesion systems that ensure robust collective migration *in vivo*.

## Results

### Integrins α3, α6 and β1b are expressed in the migrating pLLP

Among the 24 integrin heterodimers described in mammals, α3β1, α6β1 and α6β4 are dominant in epithelial tissue and α3/α6-integrins bind laminins, the major component of BMs^6^. Since the pLLP cells originate from an epithelial placode and largely maintain their epithelial properties ^35,37^, we wanted to determine whether the corresponding zebrafish paralogs (*itga3a/itga3b* and *itga6a/itga6b/itga6l)* were expressed in the pLLP during migration. Therefore, we performed *in situ* hybridization in wild-type embryos expressing the LL reporter *Tg(-0.8cldnb:lynGFP)^zf^*^106^ (hereafter: *cldnb:GFP*) ^38^. Both α3 paralogues, *itga3a* and *itga3b*, were expressed in the trailing region of the pLLP. *itga3b* exhibited broad expression in the trailing half of the pLLP throughout migration and remained expressed in deposited neuromasts (**Fig. 1A-C’’**). In contrast, *itga3a* expression was observed in small clusters along the pLLP midline, with its expression restricted to early stages of migration (24 and 30hpf) (**Fig. 1D-F’’**). While *itga6b* was expressed during the entire migration process (**Fig. 1G-I’’**), neither *itga6a* nor *itga6l* were expressed in the migrating pLLP (**Fig. S1A-B’**). In contrast to *itga3a* and *itga3b, itga6b* expression was more pronounced in the leading region, in particular towards the end of migration at 48hpf (**Fig. 1G-I’’**). Neither *itga3a/itga3b* nor *itga6b* were maternally expressed (**Fig. S1F-H**). Integrin β1 is a major dimerization partner of integrins α3 and α6 ^6^. Among the two zebrafish *itgb1* paralogues, *itgb1b* was expressed in the migrating pLLP at 30hpf but was absent at 48hpf (**Fig. 1J-L’’)** whereas *itgb1a* was not expressed in the pLLP at any stage (**Fig. S1C-E’’**), consistent with previous findings ^36^. Taken together, these findings indicate that three major epithelial integrins - *itga3a, itga3b* and *itga6b* – along with their presumptive dimerization partner *itgb1b* - are expressed in the migrating pLLP within specific domains.

**Figure 1:**
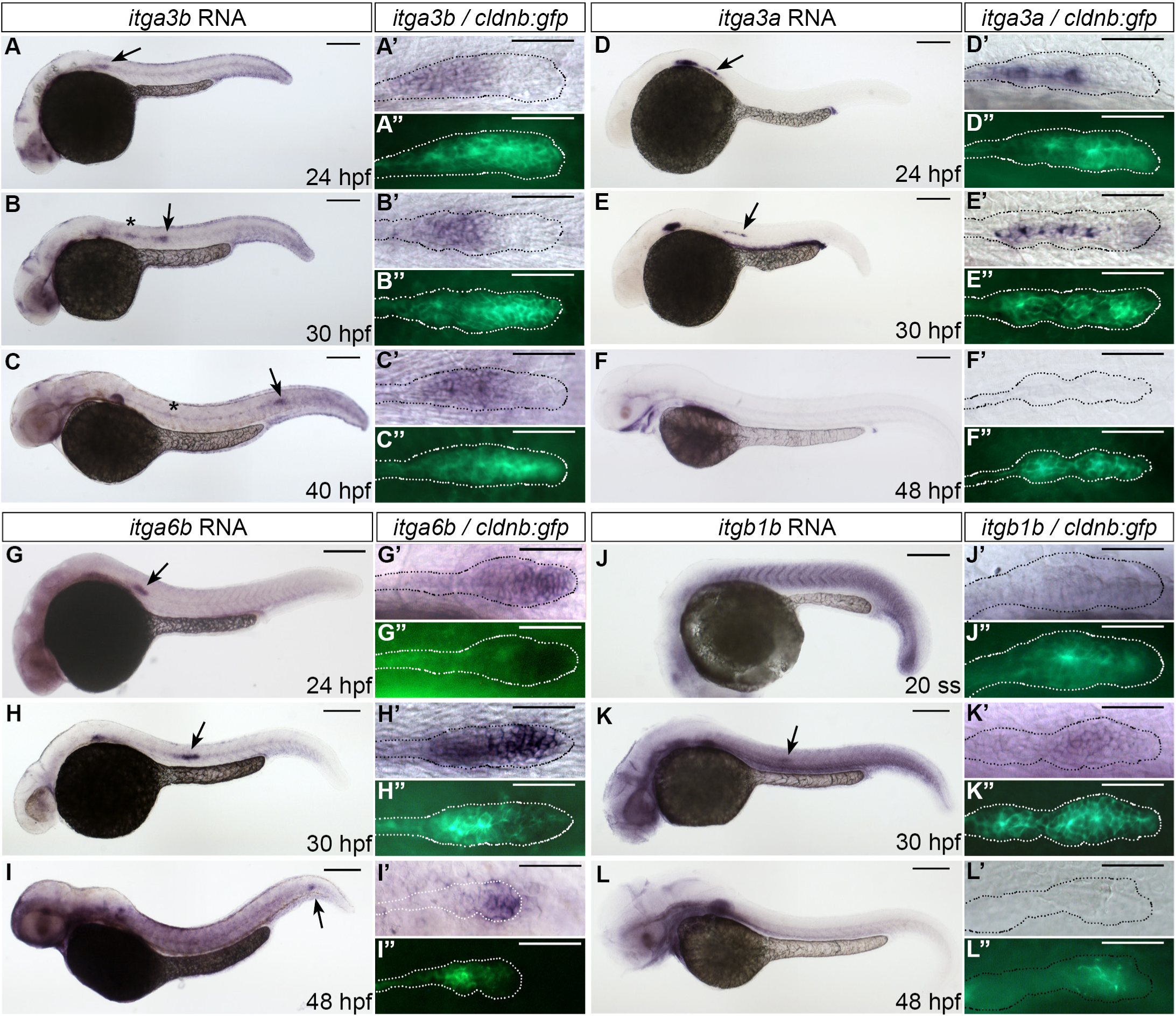
*itga3b*, *itga3a*, *itga6b* and *itgb1b* are expressed in the migrating primordium. *cldnb:GFP* embryos were stained by *in situ* hybridization (ISH) using riboprobes against *itga3b* (A-C’’), *itga3a* (D-F’’), *itga6b* (G-I’’) and *itgb1b* (J-L’’) at the indicated stages. Arrows mark the migrating pLLP. (A-L) show overview images of ISH staining. (A’-L’) show close-ups on the pLLP and (A’’-L’’) show GFP immunostaining to delineate the pLLP. Scale bars: 200 μm (A-L), 50 μm (A’-L’’).

### *itga3a, itga3b* and *itga6b* are required during pLL development

To investigate the function of these integrins in the pLLP, we first analysed the previously described *badfin (bdf)* mutants (*itga3b^fr^*^21^, Carney et al., 2010). At 40 hpf, *itga3b^−/−^*embryos exhibited no morphological defects aside from the characteristic caudal fin dysmorphogenesis ^39^ **(Fig. 2A-C, green arrow)**. Both pLLP morphology and migration appeared unaffected in *itga3b^−/−^* embryos, with no detectable defects in migration distance **(Fig. 2 A-C’’, quantification in Fig. 2J)**. Although the *bdf* mutation results only in a single amino-acid substitution (S427P), *itga3b* mRNA levels were reduced by more than 98% in maternal zygotic (MZ) *itga3b^−/−^* mutants, suggesting that the mutant transcript is degraded (**Fig. S2G**) and that *badfin* likely represents a null allele. Given that mutant mRNA degradation can trigger transcriptional adaptation ^40^, we also assessed the expression of *itga3a* and *itga6b* in *itga3b^−/−^* mutants. *itga3a* levels were reduced and *itga6b* expression remained unchanged in MZ *itga3b^−/−^* mutants (**Fig. S2G**), indicating that loss of *itga3b* is not compensated by upregulation of the expression of either its paralog *itga3a* or another laminin-binding α-integrin, *itga6b*.

**Figure 2:**
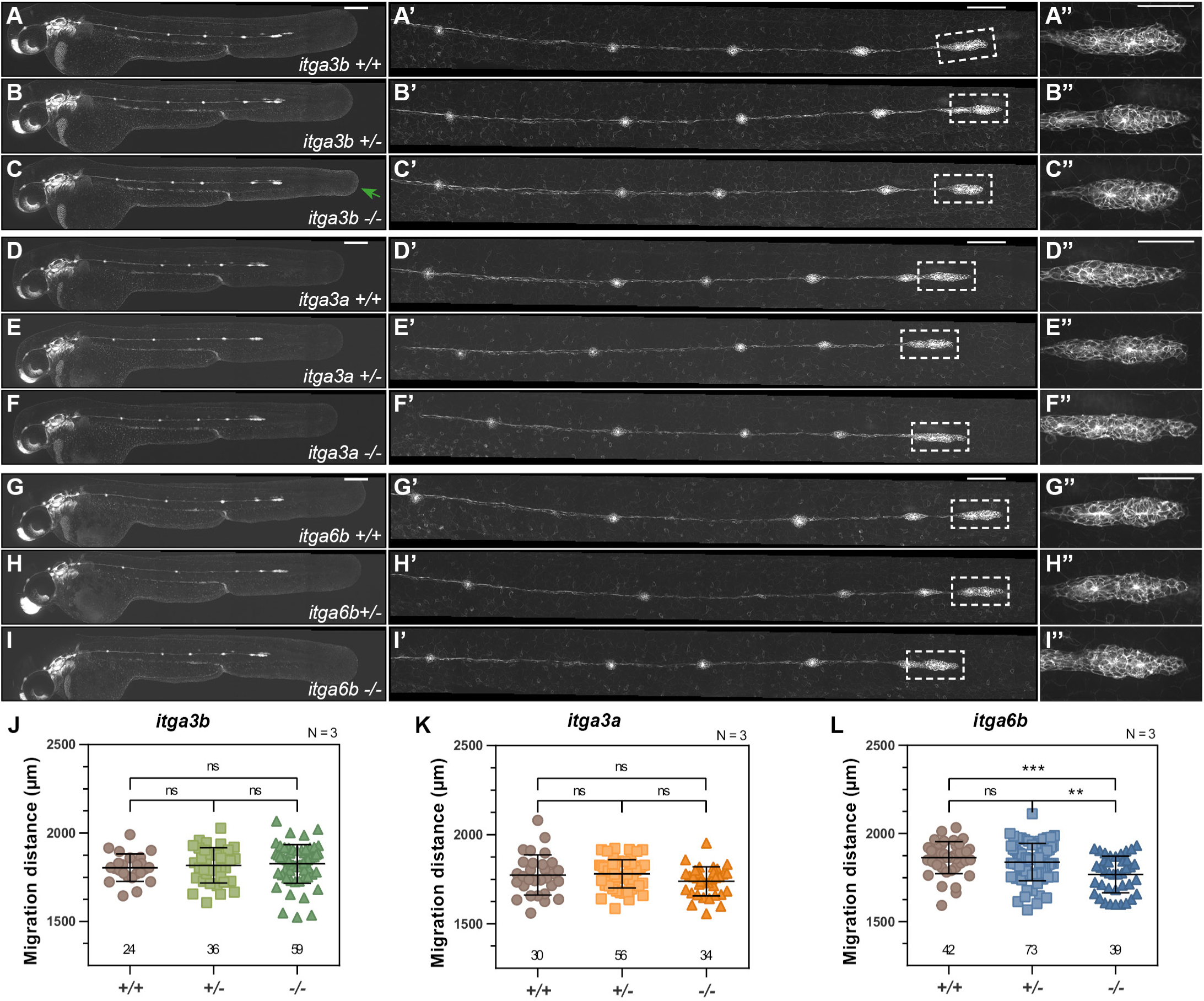
*itga6b*, but not *itga3a* or *itga3b*, is required for proper pLLP migration. (A-I’’) Representative images of *cldnb:GFP* embryos at 40 hpf of wild-type, heterozygous and homozygous *itga3b* (A-C’’)*, itga3a* (D-F’’) and *itga6b* (G-I’’) mutants, including embryo overviews (A-I), higher magnifications of the pLL (A’-I’), and a close-ups on the pLLP (A’’-I’’). A green arrow in (C) indicates the *badfin* phenotype of *itga3b* mutants. (J-L) Quantification of the distance migrated by the pLLP at 40 hpf. Statistics: One-way ANOVA with multiple comparisons; ns = non-significant; **p < 0.01; ***p < 0.001. Sample size (n) are indicated above x-axes; N = number of biological replicates. Scale bars: 200 μm (A-I), 100 μm (A’-I’) and 50 μm (A’’-I’’).

Since *itga3a* is expressed in the pLLP, it may still functionally compensate for the loss of *itga3b*. To test this, we generated *itga3a* mutants using transcription activator-like effector nucleases (TALENs), targeting the first exons shortly after the initiation codon (**Fig. S2A**). Among several recovered *itga3a* mutant alleles, we selected a 5bp deletion (allele *fu40*), which introduces a premature stop codon after 54 amino acids for further analysis (**Fig. S2B**). While *itga3a^fu^*^40^ adult fish were not viable - except for rare escapers that were drastically reduced in size (**Fig. S2C**) - *itga3a^−/−^*embryos developed normally, were morphologically indistinguishable from their siblings, and showed no defects in pLL development (**Fig. 2D-F’’, quantification in Fig. 2K**).

To assess potential functional redundancy between *itga3a* and *itga3b*, we generated *itga3a^fu^*^40^*;itga3b^fr^*^21^ double mutants. Double *itga3a^−/−^;itga3b^−/−^*embryos displayed the same *bdf* caudal fin phenotype as single *itga3b^−/−^*and were otherwise indistinguishable from their siblings **(Fig. S3, compare F to B-D)**. Despite both paralogs being expressed in the pLLP during migration, double *itga3a^−/−^;itga3b^−/−^* mutants exhibited normal pLLP morphology, migration, proneuromast formation, and deposition (**Fig. S3F,F’, quantification in Fig. S3J**). These findings suggest that integrin α3 activity is either not required in the pLL or its function is compensated by yet another integrin or an alternative protein.

To investigate the function of *itga6b,* we employed the same TALEN-based loss-of-function approach used for *itga3a* (**Fig. S2D**). We selected an allele carrying a 7bp deletion leading to a frameshift and a premature stop codon after 26 amino acids (*fu36*) (**Fig. S2E**). Similar to *itga3a^fu^*^40^, *itga6b^fu^*^36^ mutants were not viable as adults, although rare escapers with severe growth restriction were occasionally observed (**Fig. S2F**). At 40hpf, *itga6b^−/−^* embryos showed no morphological defects. However, detailed examination of the lateral line (LL) revealed a slight delay in pLLP migration **(Fig. 2G-I, quantification in Fig. 2L)**. Notably, this phenotype was subtle enough that it did not translate in a significant decrease in migration speed in 14-hour time-lapse recordings (TL) **(Fig. S3K-N, Video S5).**

We next wanted to test whether, in the absence of Itgα6b, Itgα3a and/or Itgα3b functions were required for pLLP migration and morphogenesis. For this purpose, we generated triple heterozygous *itga3a^+/-^;itga3b^+/-^; itga6b^+/-^* fish and analysed pLL formation in their offspring. Since heterozygous embryos exhibited no phenotypic abnormalities, we pooled wild-type and heterozygous individuals (indicated as “+/?”) for analysis. Single mutants derived from this lineage displayed phenotypes consistent with previous observations (**Fig. S3**). Loss of both *itga3a* and *itga6b* did not affect overall embryo morphology (**Fig. 3C-C’**) and did not enhance the mild migration delay observed in *itga6b* single mutants (Compare **Fig. 3B’ vs. 3C’, quantification in Fig. 3F**). In contrast, *itga3b^−/−^; itga6b^−/−^* double mutants (hereafter: *itga3b;6b* double mutants) exhibited a noticeably thinner caudal fin compared to the characteristic *“badfin”* of *itga3b* mutants (**Fig. 3D, green arrow**). Additionally, their pLLPs showed a significantly greater migration delay than *itga6b^−/−^* mutants, (Compare **Fig. 3D’ to 3B’ and 3A’, quantification in Fig. 3F**). A small proportion of embryos from this cross (1/64) were *itga3a^−/−^;itga3b^−/−^;itga6b^−/−^* triple mutants (hereafter: *itga3a;3b;6b* triple mutants). Additional loss of *itga3a* further enhanced the migration phenotype of *itga3b;6b* double mutants (**Fig. 3E’, 3F**). The dysmorphogenesis of the caudal fin was similar to *itga3b;6b* double mutants (**Fig. 3E, green arrow**). Despite the pronounced migration delay, both double and triple mutant pLLPs ultimately completed migration (data not shown). Time-lapse recordings from 30 hpf confirmed that both double and triple mutants demonstrated reduced migration speed from the onset and maintained this slower pace throughout the entire migration process (Fig. 3G-K, Video S1). The delayed migration of integrin double and triple mutants was accompanied by a markedly stretched pLLP morphology (**Fig. 4B-C**) compared to sibling controls (**Fig. 4A**).

**Figure 3:**
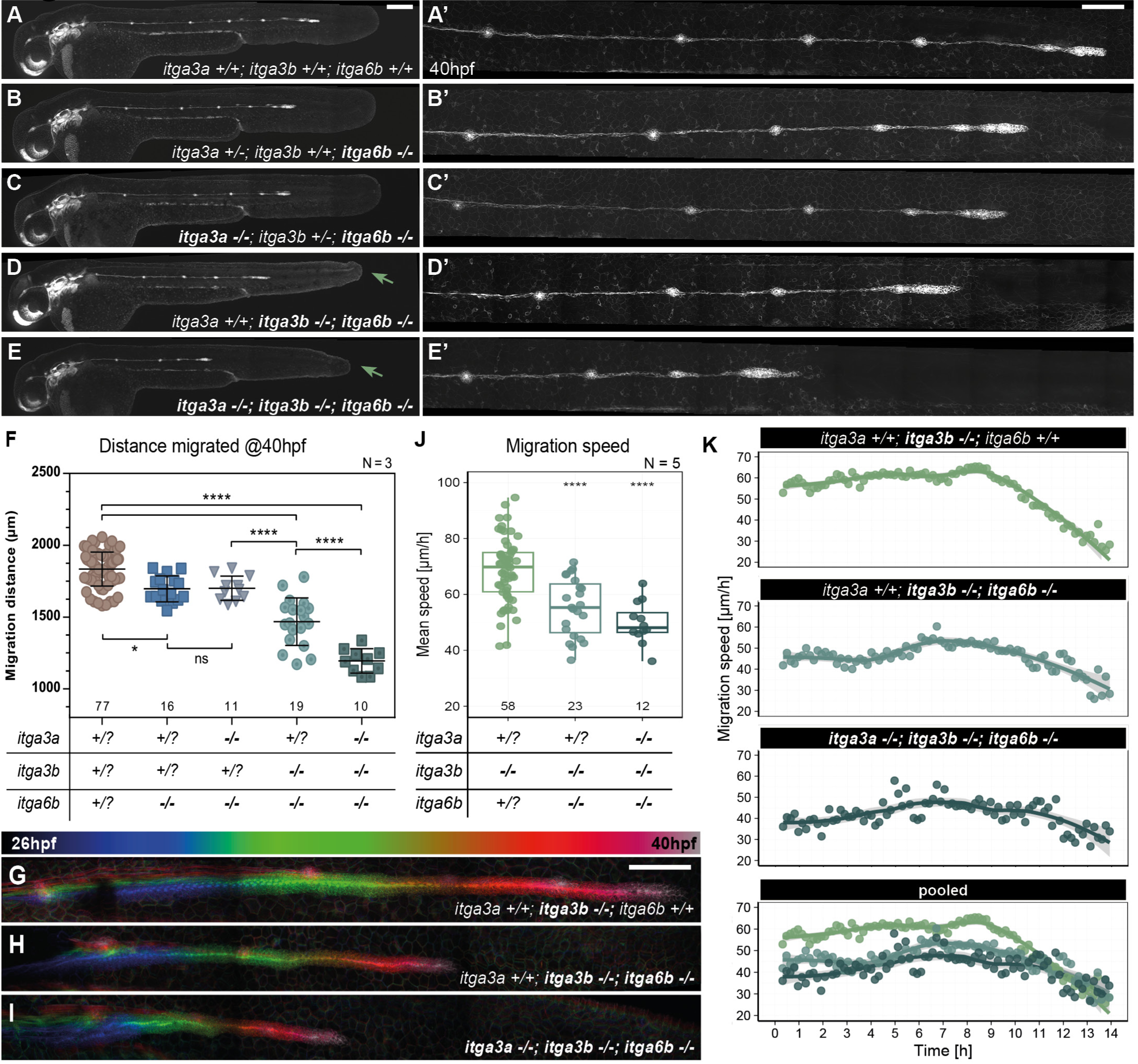
*itga3a*, *itga3b* and *itga6b* function redundantly and hierarchically to mediate pLLP migration. (A-E’) Representative images of Integrin compound mutants at 40hpf showing whole embryos (A-E) and higher magnifications of the pLL (A’-E’). Green arrows indicate the *badfin* phenotype in *itga3b;itga6b* double mutants (D) and *itga3a;itga3b;itga6b* triple mutants (E). (F) Quantification of pLLP migration distance reveals significant delays in double and triple mutants (F). (G-I) Temporally color-coded maximum intensity projection (MIPs) from 14-hour TLs of the indicated genotypes. Quantification of mean (J) and instantaneous (K) migration speed. Statistics: One-way ANOVA with multiple comparisons (F) and two-tailed Wilcoxon rank-sum test (J); ns = non-significant; *p < 0.05; ****p < 0.0001. Sample sizes (n) are indicated above x-axes. N = number of biological replicates. Scale bars: 200 µm (A-E, G-I), 100 µm (A’-E’).

**Figure 4:**
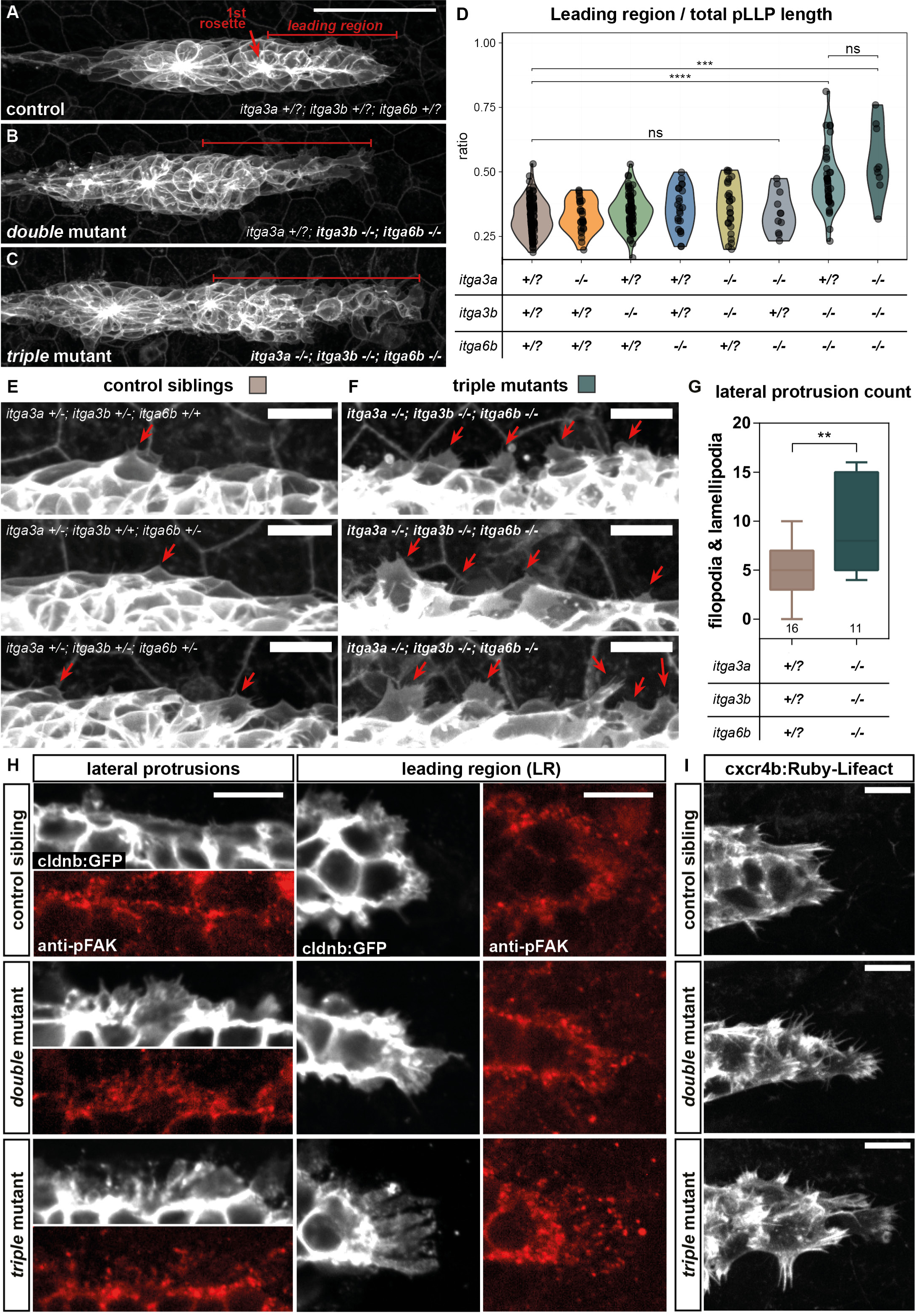
*itga3b*;*itga6b* double and *itga3a;itga3b;itga6b* triple mutant pLLPs form ectopic protrusions. (A-C) Live imaging of pLLPs in sibling control (A), *itga3b*;*itga6b* double (B) and *itga3a;itga3b;itga6b* triple (C) mutants at 40hpf. The leading region (LR) extends from the tip of the pLLP to the first rosette (red segments). (D) Quantification of the ratio LR/total pLLP length. (E-F) Close-ups showing lateral protrusions (red arrows) in mutant pLLPs and corresponding quantification (G). (H) Single z-planes of pLLP immunostained with anti-pFAK in the indicated genotypes. (I) Live imaging of *cxcr4b:R-Lifeact* embryos reveals ectopic leading protrusions in mutants. Statistics: Pairwise Wilcoxon rank-sum test with Holm-adjusted p-values (D) and two-tailed Wilcoxon rank-sum test (G). ns = non-significant; **p < 0.01; ***p < 0.001; ****p < 0.0001. Scale bars: 50 μm (A-C), 10 μm (E-F, H-I).

Specifically, the proneuromast-free leading region (LR) was significantly more extended in *itga3b;6b* double and *itga3a;3b;6b* triple mutants compared to their siblings (**Fig. 4D**). Imaging of the live actin reporter *Tg(cxcr4b:Ruby-Lifeact)* (hereafter: *cxcr4b:R-Lifeact*) demonstrated actin-based filopodia and lamellipodia within extensively stretched LRs (**Fig. 4I)**. To minimize bias due to the strong phenotype of the LR, we quantified lateral protrusions in blinded images (**Fig. 4E-F**) and found a significant increase in the number of lateral protrusions in triple integrin mutants compared to their control siblings (**Fig. 4G**). Given that integrins are core components of focal adhesions (FAs), we hypothesized that the increased protrusive activity in *itga3b;6b* double and *itga3a;3b;6b* triple mutants might result from impaired FA stabilization. To investigate this, we examined the localization of phosphorylated focal-adhesion kinase (pFAK), a marker for active FAs ^41,42^. In control siblings, pFAK localized in clusters at the periphery of the pLLP, particularly within cell protrusions **(Fig. 4H)**. Similarly, pFAK was detected within the extensive lamellipodia and filopodia of *itga3b;6b* double and *itga3a;3b;6b* triple mutants, indicating that despite the increased protrusive activity, FA formed normally in these mutants.

In summary, our analysis revealed a front-to-rear polarized expression of adhesion receptors in the pLLP with *itga6b* in the leading and *itga3a/itga3b* in the trailing region. Only the loss of *itga6b* resulted in a mild delay in migration, while the loss of *itga3a* or *itga3b*, either individually or in combination, had no detectable impact on pLLP migration or morphology. Itga3b only became essential in the absence of Itga6b and Itga3a only in the absence of both Itga3b and Itga6b. These findings highlight a highly robust and partially redundant adhesion machinery with an increasing hierarchical requirement for Itga3a, Itga3b, and ultimately Itga6b during posterior lateral line morphogenesis.

### Lama5 is part of the basement membrane upon which pLLP cells migrate and is necessary for its integrity

Integrins mediate cell adhesion to the extracellular matrix and are classified based on their ligand affinity ^6^. Since α3 and α6 integrins primarily bind laminins, we first confirmed the presence of laminins around the pLLP by performing pan-laminin immunostaining on whole embryos (**Fig. 5A-A’**) and cryosections (**Fig. 5B-B’**). Next to the somite boundaries (**Fig. 5A-B, yellow arrows**), the antibody stained a basement membrane (BM)-like sheet beneath the pLLP (**Fig. 5A’-B’, white arrows**). Laminins were also detected at pLLP cell membranes and at the rosette centers, though at lower levels (**Fig. 5B-B’, asterisk**). This distribution aligns with the reported localization of the core BM protein Lamc1 ^36^. To identify the specific laminin ligand involved in pLLP migration, we focused on laminin-α5 (Lama5), a key α-chain component of BM underlying epithelial tissues, and known to interact with α3- and α6-integrins ^6,43,44^. To determine whether Lama5 is present in the BM supporting pLLP migration, we performed immunostaining using an antibody against the mouse Lama5 protein (generous gift from the Sorokin lab, Hannocks *et al.*, 2018). in the recently established *TgBAC(lamc1:lamc1-sfGFP)^sk^*^116^*^Tg^* BAC reporter line (hereafter: *lamc1-GFP)* ^36^. Lama5 staining was detected in the BM beneath the pLLP (**Fig. 5C, white arrow**), where it co-localized with Lamc1-GFP (**Fig. 5C’-C’’**). However, in contrast to Lamc1-GFP, Lama5 was absent from the somite boundaries (**Fig. 5C’’, yellow arrow**). This result indicated that Lama5 is one of the Laminin α subunit present in the epidermal BM over which the pLLP migrates and may function as a ligand, together with Lamc1, for integrin receptors expressed in the pLLP, such as Itgα3a/b or Itgα6b.

**Figure 5:**
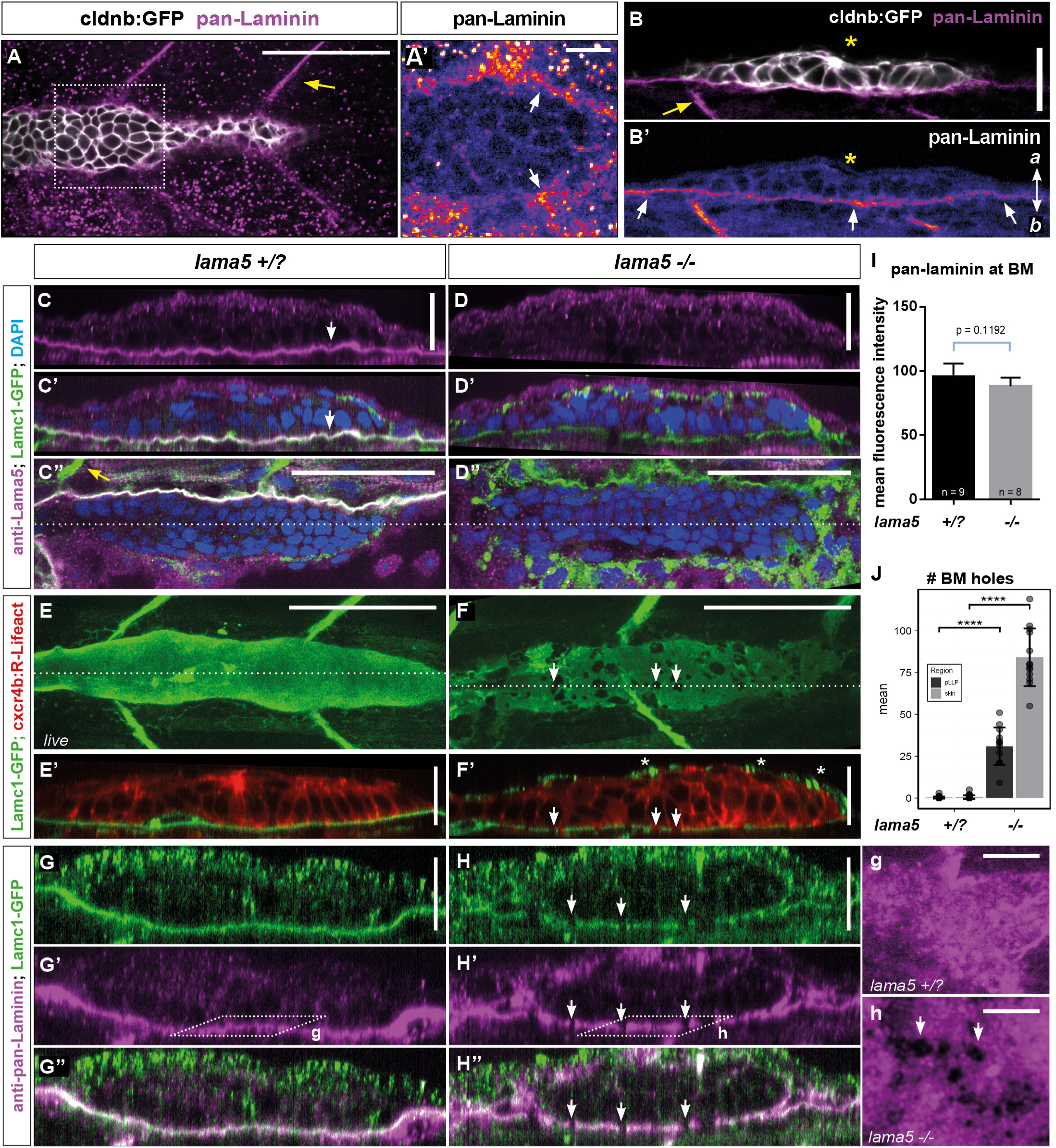
Laminin α5 is an integral component of the epidermal BM. (A-B’) Pan-laminin immunostaining in whole-mount embryos (A-A’) and transverse cryosections (B-B’). Yellow arrows point to somite boundaries (A, B); white arrows mark the epidermal BM (A’, B’); white asterisk marks laminin enrichment at the center of epithelial rosettes (B-B’). (C-D’’) Anti-Lama5 immunostaining in *lama5^+/+^* or *lama5^+/-^* (*lama5^+/?^*) (C-C’’) and *lama5^−/−^* (D-D’’) embryos carrying the *lamc1-GFP* transgene. Orthogonal views (C-D’) and single z-planes (C’’,D’’) show Lama5 and Lamc1 in the epidermal BM (white arrow in C-C’) in controls. Lama5, but not Lamc1, is missing in the BM in *lama5* mutants (D-D’’). (E-F’) Live imaging of *cxcr4b:R-Lifeact* (red) and *lamc1-GFP* (green) highlights BM holes in *lama5* mutants (Arrows). Shown are maximum intensity projections (MIPs) of 5µm-substacks (E-F) and single orthogonal views (E’-F’). (J) Quantified total number of holes below pLLP and adjacent skin. (G-H’’) Pan-laminin immunostaining in *lamc1-GFP* embryos. Orthogonal views (G-H’’) and close-ups of single z-planes (g, h) as indicated in (G’, H’) show similar discontinuities in the BM in *lama5* mutants (white arrows in H-H’’, h). (I) Quantification of pan-laminin signal intensity. Statistics: Two-tailed Wilcoxon rank-sum test. ns = non-significant; ****p < 0.0001. Scale bars 50 µm for top-views (A, C’’-D’’, E-F), 20 µm for optical sections (B-B’, C-D’, E’-F’, G-H’’) and 10 µm for close-ups (A’, g-h). a = apical, b = basal.

To investigate Lama5 function during pLL morphogenesis, we used the previously described *fransen* mutant (allele *tc17,* Carney *et al.*, 2010). Lama5 immunostaining was absent from the epidermal BM in *lama5* mutants **(Fig. 5D)**, while the Lamc1-GFP signal remained present **(Fig. 5D’-D’’)**. This confirms the antibody’s specificity for zebrafish Lama5, and that the truncated protein produced by the *tc17* nonsense mutation is either degraded or not secreted. Interestingly, Lamc1-GFP formed aggregates within the basal epidermal cells of *lama5^−/−^* mutants **(Fig. 5F’, white asterisks)**, suggesting that Lamc1 is not properly secreted in the absence of Lama5. This further implies that Lama5 is required for the proper assembly or export of Lamc1, supporting the idea that both proteins are components of the same Laminin trimer. Additionally, the BM labelled by Lamc1-GFP appeared discontinuous in *lama5* mutants (**Fig. 5E-H’’**). Live imaging confirmed the presence of holes within the otherwise continuous, sheet-like Lamc1-GFP signal **(compare Fig. 5E and 5F)**. Quantification of these gaps beneath the pLLP and in the surrounding BM indicated that this phenotype was not restricted to the pLLP migration path (**Fig. 5J**). Pan-laminin immunostaining confirmed the presence of these gaps, ruling out the possibility that other laminin isoforms are present in these gaps (**Fig. 5H-H’, 5g-h**). Notably, overall pan-laminin staining intensity remained unchanged (**Fig. 5g-h, quantification in Fig. 5I**).

To gain ultrastructural insight into the BM defects, we performed transmission electron microscopy (TEM) at 32 hpf. In control embryos, the epidermal BM was approximately 40nm thick and contained irregularly spaced adepidermal granules (**Fig. S4A-A’**), consistent with previous reports that describe granule appearance between 24 and 48 hpf ^46^. In contrast, *lama5* mutants lacked adepidermal granules in BM regions of wild-type thickness (**Fig. S4B**). These regions were rare, as the BM was frequently compromised by expanded, bloated segments, and in some areas, the lamina densa was no longer discernible (**Fig. S4C**). We believe these areas correspond to the gaps observed by live imaging of Lamc1-GFP, suggesting that the discontinuities observed with the light microscope reflect underlying structural defects in BM organization.

Taken together, these results indicate that Lama5 is present beneath the pLLP and is required for proper Lamc1 secretion and the assembly of an intact BM. The structural discontinuities observed by live imaging were corroborated by TEM, which revealed bloated BM segments and loss of lamina densa in *lama5* mutants. Furthermore, the persistence of Lamc1-GFP and pan-laminin signal in *lama5^−/−^* mutants implies that at least one other laminin α-subunit forms a trimer and is secreted together with Lamc1 into the epidermal BM.

### Lama5 constitutes a substrate for the posterior lateral line but is not required for pLLP migration

Given that Lama5 is present beneath the migrating pLLP and that the *lama5* mutation disrupts BM organization, we next investigated the consequences of *lama5* loss on pLL formation. In addition to the previously reported caudal fin dysmorphogenesis ^39^ **(Fig. 6A-D, red arrows in 6B, D)**, we observed several striking abnormalities in the pLL. During migration, the pLLP deposits neuromasts, which are connected by a chain of interneuromast cells (INCs) (**Fig. 6A’,a,C’,c**) ^47^. In *lama5^+/?^* embryos, INCs organized into a smooth, stable, single-layered structure surrounding the pLL nerve **(Fig. 6a, 6c and Video S2)**. In contrast, INCs in *lama5^−/−^* mutants were highly destabilised, exhibiting increased protrusive and migratory activity **(Fig. 6B’, b, Video S3).** This led to the formation of large gaps within the chain, leaving segments of the pLL nerve exposed (**Video S4**) **(Fig. 6B,b,D,d)**. Quantification of these defects at 40hpf revealed that, on average, 10% of the total INC chain is interrupted by gaps in *lama5^−/−^* mutants **(Fig. 6E)**.

**Figure 6:**
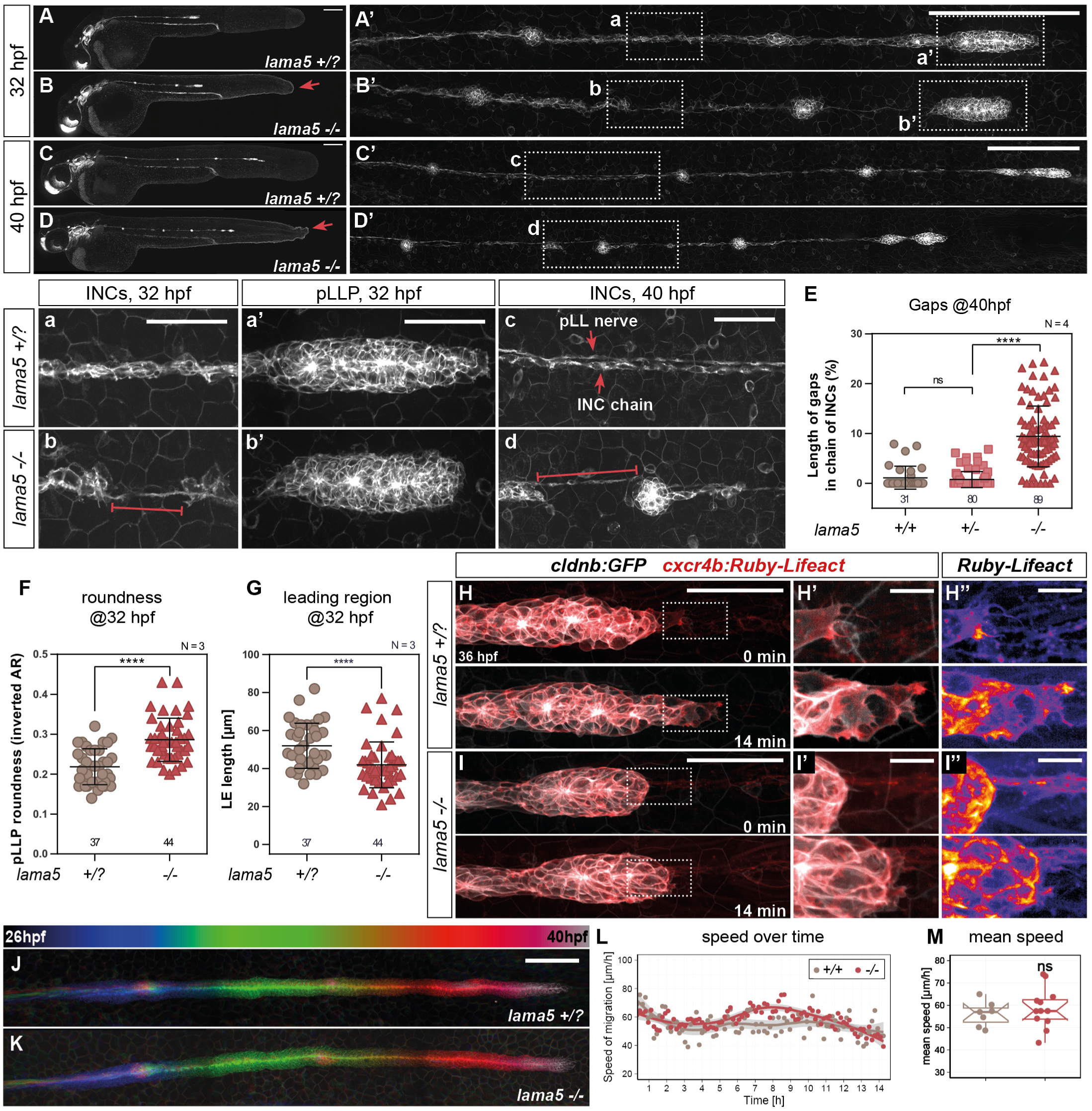
Laminin α5 is required as a substrate for LL morphogenesis. (A-D’) Representative images of *lama5^+/?^* and *lama5*^−/−^ *cldnb:gfp* embryos fixed at 32hpf and 40hpf showing whole embryos (A-D) and higher magnifications of the pLL (A’-D’). Red arrows in B, D point to the characteristic “*fransen”* phenotype in the caudal fin of *lama5* mutants. (a-d) Close-ups as indicated in A’-D’ showing the disrupted INC chain in *lama5^−/−^* embryos at 32 hpf (a-b) and 40 hpf (c-d). (E) Quantification of INC gap length relative to total LL length at 40hpf. (a’-b’) Close-ups as indicated in (A’-D’) showing that *lama5* mutant pLLPs are rounder (F) with a reduced leading region (G). (H-I’’) Snapshots of a time-lapse using the actin reporter *cxcr4b:R-Lifeact.* (H’-I’) close-ups of the leading cells corresponding to the boxes in (H-I). (H’’-I’’) Fire LUT applied to red channel highlights actin accumulation in the leading cells. (J-M) 14-hour-TL shows that pLLP migration speed is not significantly lower in *lama5* mutants. (J-K) Temporally color-coded MIPs shows migration over time. (L-M) Quantification of instantaneous (L) and average (M) speeds. Statistics: One-way ANOVA without (F-G) or with multiple comparisons (E); two-tailed Wilcoxon rank-sum test (M). ns = non-significant; ****p < 0.0001. Scale bars: 50 μm (A), 10 μm (E-F, H-I).

Furthermore, *lama5^−/−^* pLLPs were significantly rounder than those of control siblings throughout migration (**Fig. 6a’-b’, H-I, quantification in 6F, S4E**). This increased roundness coincided with a reduced leading region (LR) in *lama5* mutant primordia (**Fig. 6G**). Using the *cxcr4b:R-Lifeact* reporter, we qualitatively compared actin-based protrusions in the LR but could not observe any obvious difference (**Fig. 6H-I’’**). Surprisingly, despite pLLP rounding, LR shortening, and a discontinuous BM lacking Lama5, *lama5^−/−^* primordia migrated at speeds comparable to those of control siblings (**Fig. 6J-K, quantification in 6L-M, Video S5**). Only a slight reduction in total migration distance at 40 hpf was detected (**Fig. S4D**), similar to *itga6b* mutants (**Fig. 2L**).

In summary, loss of Lama5 differentially affected the deposited LL cells and the migrating pLLP. While INCs were destabilized and hypermotile in the absence of Lama5, *lama5* mutant primordia were notably rounder with a reduced leading region, yet overall migration remained largely normal. The rounding phenotype contrasts sharply with the elongated pLLPs observed in double and triple integrin mutants. Since *lama5* loss did not phenocopy integrin mutants, we hypothesize that Lama5 likely interacts with an alternative receptor expressed in the LR. Conversely, Lama5 is unlikely to serve as a unique ligand for integrins α3 and α6 in the pLLP.

### Genetic interactions between *lama5* and *itga3/a6b* in pLL development

To test potential genetic interactions between laminin α5 and integrin α3 and α6b, we generated double and triple mutants, and analysed the phenotype of their progeny. The loss of either Itga3a or Itga3b did not enhance the *lama5^−/−^* phenotype (**Fig. S5E, H-I, Q**). In contrast, the combined loss of Itga6b receptor and Lama5 ligand had a strong impact on the pLL development. Compared to the mild migration delay observed in single *lama5* mutants (**Fig. 7A-C**), the additional loss of one copy of *itga6b* resulted in a significant reduction in migration distance (**Fig. 7D, quantification in F**). At 40hpf, these primordia had only migrated as far as the end of the yolk extension (**Fig. 7D)**. Despite this delay, these primordia successfully completed migration and reached the tip of the fin (data not shown). Strikingly, the additional loss of the second *itga6b* copy in *lama5^−/−^;itga6b^−/−^* double mutants further worsened the migration defect with the pLLP reaching only the middle of the yolk extension at 40hpf (**Fig. 7E**). Most notably, the pLLP of *lama5^−/−^;itga6b^−/−^*mutants stalled, failing to migrate past the end of the yolk extension even at 5 dpf (data not shown). The additional loss of either *itga3a* or *itga3b* did not enhance the *lama5^−/−^;itga6b^−/−^* migration phenotype **(Fig. S5 O-P)**, nor did the depletion of both *itga3a* and *itga3b* enhance the *lama5^−/−^; itga6^+/-^* migration phenotype **(Fig. S5N compare to S5K)**. Quadruple mutants could not be obtained in this study.

**Figure 7:**
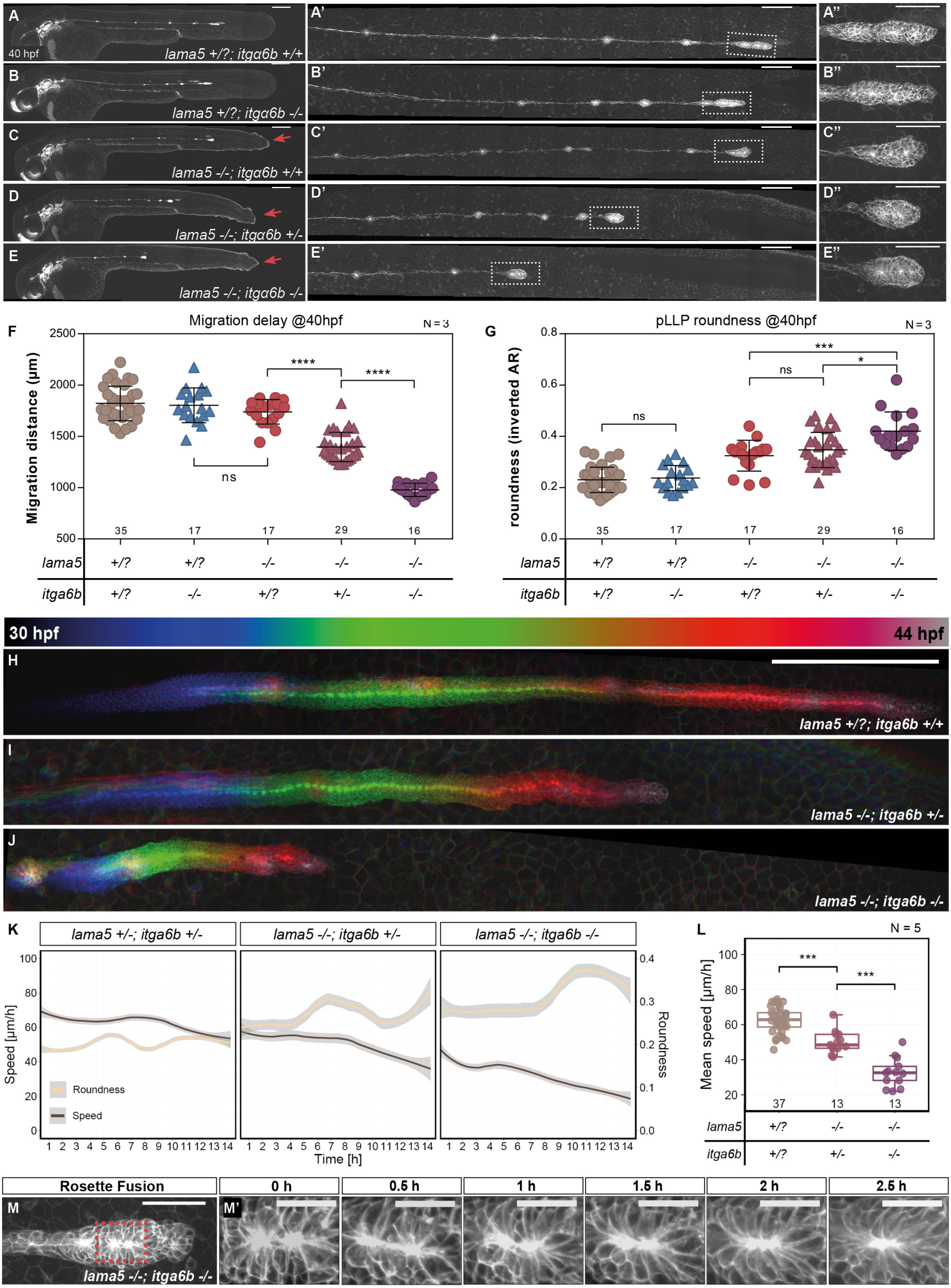
*lama5* and *itga6b* interact genetically to maintain pLLP migration. (A-E’’) Representative images of *lama5;itga6b* compound mutant embryos carrying the *cldnb:gfp* transgene fixed at 40hpf. Compared to controls (A-A’) and single mutants (B-C’), pLLP migration is impaired in *lama5^−/−^;itga6b^+/-^* (D-D’) and even more severely in *lama5^−/−^ ;itga6b^−/−^* double mutants (E-E’). (F) Quantification of the distance migrated by the pLLP at 40hpf. (A’’-E’’) Close-ups of the pLLP show increased roundness in compounds mutants; quantification shown in (G). (H-J) Temporally color-coded MIPs of 14-hour time-lapse imaging of sibling control vs. *lama5;itga6b* compound mutants with the indicated genotype. (K) Quantification of instantaneous speed and pLLP roundness over time. (L) Quantification of the mean migration speed. (M-M’) Example of a rosette fusion event in a *lama5^−/−^;itga6b^−/−^* double mutant pLLP. Sample sizes (n) are indicated above x-axes. N is the number of biological replicates. Statistics: One-way ANOVA with multiple comparisons (F-G); Pairwise Wilcoxon rank-sum test (L). ns = non-significant; * p < 0.05; *** p < 0.001;**** p < 0.0001. Scale bars: 200 µm (A-E, H-J), 100 µm (A’-E’), 50 µm (A’’-E’’, M), 10 µm (M’).

Fourteen-hour time-lapse recordings starting at 30 hpf further revealed that *lama5^−/−^; itga6b^−/−^* primordia exhibited a reduced migration speed from the onset **(Fig. 7K, Video S5)**. While wild-type and single mutant primordia migrated at an average speed of 60 µm/h **(Fig. 7L, S6G)**, *lama5^−/−^; itga6b^−/−^* mutants exhibited a markedly reduced mean speed of 35 µm/h **(Fig. 7L)**. Initially migrating at about 40 µm/h, these primordia progressively slowed to below 20 µm/h by the end of the time-lapse **(Fig. 7K)**, accounting for the gradual stalling phenotype observed in the later stages of migration **(Video S5)**. Consistent with the migration distance analysis, *lama5^−/−^;itga6b^+/-^*mutants displayed an intermediate phenotype, with migration speeds slightly reduced compared to control groups but significantly higher than *lama5^−/−^;itga6b^−/−^*double mutants **(Fig. 7K-L, Video S5)**.

The migration defect observed in *lama5;itga6b* mutant primordia was associated with an extremely rounded morphology **(Fig. 7D’’-E’’)**, more severe than that observed in *lama5* single mutants (**Fig. 7G**). This phenotype was also evident in the time-lapse recordings (**Fig. 7K**). Despite their migration impairment and extreme rounding, *lama5;itga6b* mutant primordia retained the ability to dynamically extend and retract protrusions in the direction of migration (**Fig S6 I-I’ + Video S6**). In some instances, the leading edge appeared to split, as if bypassing an obstacle (**Fig S6 C + Video S7**). In other instances, we observed rosette centres fusing, presumably due to the tissue pushing from the rear while failing to stretch in the direction of migration (**Fig. 7 M-M’,**). Considering that the rear appears to be pushing and the leading cells are actively protruding, the inability to sustain migration remains puzzling. To our knowledge, such a stalling phenotype of the pLLP has not been previously reported.

In summary, our findings demonstrate that the combined loss of *lama5* and *itga6b* results in severely impaired migration and extensive tissue rounding. Through a systematic analysis of receptor and receptor-ligand compound mutants, we show that *lama5* and *itga6b* interact genetically, further supporting their functional interplay in pLLP migration.

## Discussion

In the present study, we provide a comprehensive analysis of the Integrin-Laminin adhesion machinery in a collectively migrating epithelium *in vivo*. We demonstrate that the pLLP uses a front-rear polarized adhesion machinery involving α3 and α6 integrin subunits and laminin α5 as a key extracellular substrate. Through systematic genetic analysis, we reveal partial redundancy and a hierarchical requirement among integrins and identify a strong genetic interaction between *lama5* and the hierarchically dominant integrin subunit *itga6b*. While combined loss of integrins causes overengagement of leading cells and decreased overall tissue speed, simultaneous depletion of *lama5* and *itga6b* results in severely diminished leading cell motility, completely stalling migration. These findings uncover a dual adhesion system that ensures robust collective migration and provide general principles of epithelial tissue dynamics *in vivo*.

### Composition of the pLLP migration substrate

We show here for the first time that laminin α5 is an essential component of the epidermal BM supporting pLLP migration. In *lama5* mutants, Lamc1-GFP accumulates in epidermal cells, suggesting that Lama5 is required for proper secretion of Lamc1. This aligns with findings by Yurchenko et al., who showed that secretion of laminin β/γ-chains requires co-expression of an α-chain, as laminins are secreted as pre-assembled heterotrimers ^48^. In mammals, the α5-chain has been shown to trimerize with β1 or β2 and γ1-3 chains ^49^. In zebrafish, *lamb1a*, *lamb4* and *lamc1* are expressed in the embryonic epidermis ^50–53^, consistent with the presence of laminin-511, one of the most prominent epithelial BM laminins, in the epidermal BM during pLLP migration.

*lama5* mutants displayed holes in the otherwise sheet-like BM labelled by Lamc1-GFP. This is consistent with discontinuous BM formation in *Lama5*-deficient murine skin and dental epithelia ^17,18,54^. Similarly, Webb *et al.* reported large clusters of laminin immunoreactivity in the epidermis of *lama5* mutant zebrafish and interpreted this as evidence of a discontinuous BM ^55^. Altogether these observations support the essential role of Lama5 in establishing intact epithelial basement membranes.

The presence of a discontinuous BM suggests partial compensation by other γ1-containing trimers. In mice, loss of Laminin α5 is partially compensated by upregulation of Laminin α1, α2, or α4 chains ^56^, and similar compensatory mechanisms may exist in zebrafish. Additionally, recent work shows that *lama2* overexpression in muscle rescues myelination defects in LL Schwann cells in zebrafish, suggesting that the underlying muscle may contribute to the epidermal BM ^57^. Further investigation will be necessary to elucidate the physiological composition and origin of the embryonic epidermal BM, the substrate for the migrating pLLP cells, and its compensatory mechanisms in *lama5* mutants.

### Dual function of Laminin α5 in the lateral line

Our findings reveal opposing functions for Lama5 in the context of LL development: (i) promoting pLLP leading edge extension and (ii) inhibiting INC migration. Essentially, Lama5 seems to both drive and restrict migratory behavior. One possible explanation is the polarized receptor expression, with *itga6b* enriched in the leading/middle region, while *itga3b* is restricted to the trailing region and maintained in the INC and deposited neuromasts. Such spatially regulated integrin expression resembles “integrin switching,” a mechanism known to modulate cell adhesivity and migration ^58–60^.

ECM remodeling may also contribute to this dual function. ECM remodeling is a well-established process during metastasis where tumor cells acquire the capability to break the BM and invade other tissues ^61^. Proteolytic cleavage of laminin-511 by MT1-MMP/MMP14 promotes cancer cell migration and angiogenesis *in vitro* ^62,63^. It is therefore conceivable that the BM underneath the pLLP is similarly remodeled as the pLLP migrates. Remarkably, MT1-MMP and integrins exhibit mutual regulation which is associated with tumor progression - including stabilization of MT1-MMP at the cell membrane via integrin binding and, conversely, proteolytic cleavage of integrins by MT1-MMP ^64^. Polarized receptor expression, as discussed above, could thus contribute to localized ECM remodeling.

Alternatively, the effect of Lama5 on the INC may be indirect via the epidermis: weakening epidermis–BM adhesion in *lama5* mutants could account for the hyperprotrusive activity of the INC leading to its disruption. In contrast, the effect of Lama5 on the pLLP would be direct via receptors expressed in pLLP cells supporting migration and extension of the leading region.

### Polarized integrin expression coordinates migration

Our systematic genetic analysis uncovered strikingly opposite phenotypes: dramatic rounding in *lama5;itga6b* mutants versus stretched and protrusive pLLPs in double and triple integrin mutants. Single *lama5* mutants exhibited a rounder pLLP with a reduced leading edge, suggesting that a receptor in the leading region binds Lama5. The fact that none of the *itga3a*, *itga3b* or *itga6b* mutants (single, double or triple) phenocopied the rounding, but instead even led to a stretched and more protrusive leading region, supports the existence of yet another receptor for Lama5 in the leading region. Itgβ1 is the most ubiquitous β-subunit and its loss leads to rounding of the pLLP ^36^. This suggests that Lama5 engages another β1-containing integrin, which is not α3β1 nor α6β1, in the leading region. Integrin α7β1, classically associated with Laminin α2 in muscle tissue, was shown to also bind Lama5, making it an intriguing candidate ^65,66^.

In the trailing region, Itga3a and Itga3b are candidate receptors for Lama5, as the additional loss of *itga3a/itga3b* did not worsen the *lama5* phenotype. However, complete loss of Itga3 function indicates that Itga3-Lama5 interaction is neither required for migration nor for INC stabilization. This suggests that another receptor may compensate for the absence of Itga3 in the trailing region.

In contrast, additional loss of *itga6b* led to a significant increase of the *lama5* rounding phenotype, a severe decrease in pLLP migration speed and ultimately a complete stop of migration. This genetic interaction indicates that Itga6b largely compensates for the absence of Lama5 via binding to unknown ECM component(s) in the leading region. Thus, Lama5 is not, or not the only ligand for Itga6b. Conversely, Itga6b cannot be the main Lama5 receptor. The severe rounding and migration phenotype is consistent with the enrichment of *itga6b* in the leading region and also with a growing number of studies showing a non-canonical role for Itga6b, beyond the formation of hemidesmosomes, in cell migration, particularly of invasive cells in different types of cancer ^67–70^.

### Compatibility with non-canonical “rear-pushing model”

The classic model of cell migration emphasizes traction forces arising at the leading edge of cells and cell collectives ^10^. However, rear-driven mechanisms have also been observed. For instance, Zebrafish and *Xenopus* cranial neural crest cell migration is driven by a supracellular actomyosin cable at the back of the cluster ^71^. Further, using an elegant, *in vivo* approach for traction force microscopy, Yamaguchi et al. recently demonstrated that the pLLP exerts highest stresses in the middle/back, essentially pushing the tissue forward with a rear motor as opposed to pulling at the front ^36^.

Our finding that the reduced leading region in *lama5* mutant pLLP does not impair overall migration supports the idea that the movement is primarily driven by pushing forces from the rear. Furthermore, the extensive stretching of the leading region in double and triple integrin mutants may be a consequence of this mechanism. Indeed, reduced pushing forces from the rear might lead to overengagement of the leading cells as a compensation mechanism. This is consistent with other contexts, such as loss of Cxcr7 or Fgf activity, in which the pLLP trailing region fails to migrate effectively and the leading region extends abnormally ^35,72,73^. Additionally, the ability of *lama5;itga6b* double mutant pLLPs to initiate migration in the first place indicates that the rear pushing machinery remains principally functional. Moreover, we observed fusion of adjacent epithelial rosettes in these mutants, likely due to the trailing cells continuing to push while the leading region fails to extend ^74^. As *lama5;itga6b* double mutant pLLPs progressively round up and slow down until migration stalls completely, we propose that extension of the leading region is required to direct rear-generated pushing forces into forward motion.

In conclusion, we uncovered here two partially redundant cell-substrate adhesion systems that together ensure robustness and coordination of collective epithelial migration. One is laminin α5-dependent, engaging an as-yet unidentified receptor in the leading region and possibly integrin α3a/b in the trailing region. The other relies on integrin α6b in the leading region, which binds to another ECM ligand, and largely compensates for the loss of Laminin α5. Building on the concept of rear-driven migration, we propose that the pLLP uses a multilevel integrin machinery with a pushing motor at the rear that generates force, and leading cells that channel and translate this force into forward movement. These results establish fundamental principles of epithelial adhesion and collective migration with implications for development, tissue homeostasis, and pathological processes such as cancer invasion.

## Acknowledgments

We thank Dr. Holger Knaut for sharing the *Lamc1-GFP* transgenic line and Dr. Lydia Sorokin for kindly providing the anti-Lama5 antibody. We are grateful to the animal caretaker team for excellent fish care, to M. Kamprad, A. Niedztwezki and M. Heyde for technical assistance, and to our bachelor and master students V. Meckel, V. Tkalec and T. Urosevic for their help. We thank S. Eimer for the usage of the electron microscopy facility and critically reading the manuscript. We thank all members of the Lecaudey Lab for helpful discussions and support throughout the project. This work was supported by the Deutsche Forschungsgemeinschaft (German Research Council) [DFG-GRK1104 to V.L., DFG-EXC294 (BIOSS Centre for Biological Signalling Studies) to V.L, DFG-SFB850-A2 to V.L., INST 161/896-1 to V.L. and SPP1782 LE2681 to V.L.]

## Author Contributions

A.M. and N.M designed and performed most of the experiments and analyzed the data. P.A.K. and M.B. performed some experiments. C.D. generated the *itga6b* mutant line and performed some initial experiments. A.M, N.M. and V.L. conceived the project, interpreted the data and wrote the manuscript. V.L. acquired funding, coordinated the project and supervised the work.

## Supplementary Figure Legends

**S1:**
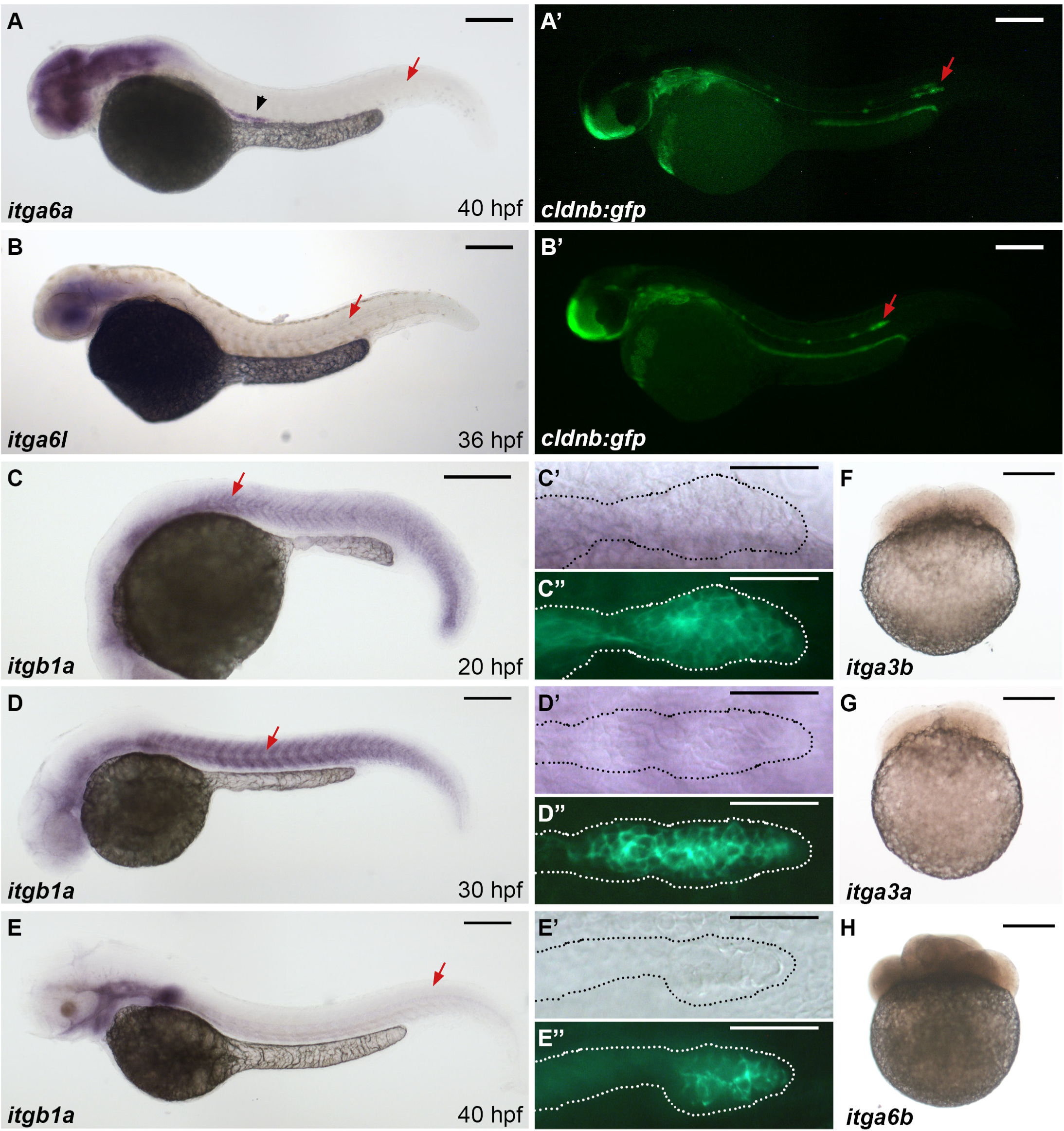
Integrin expression by ISH. (A-E’’) *itga6a*, *itga6l* and *itgb1a* are not expressed in the migrating pLLP at the indicated stages. (F-H) *itga3a*, *itga3b* and *itga6b* mRNA are not provided maternally. Scale bars: 200µm (A-E), 50µm (C’-E’’, F-H).

**S2:**
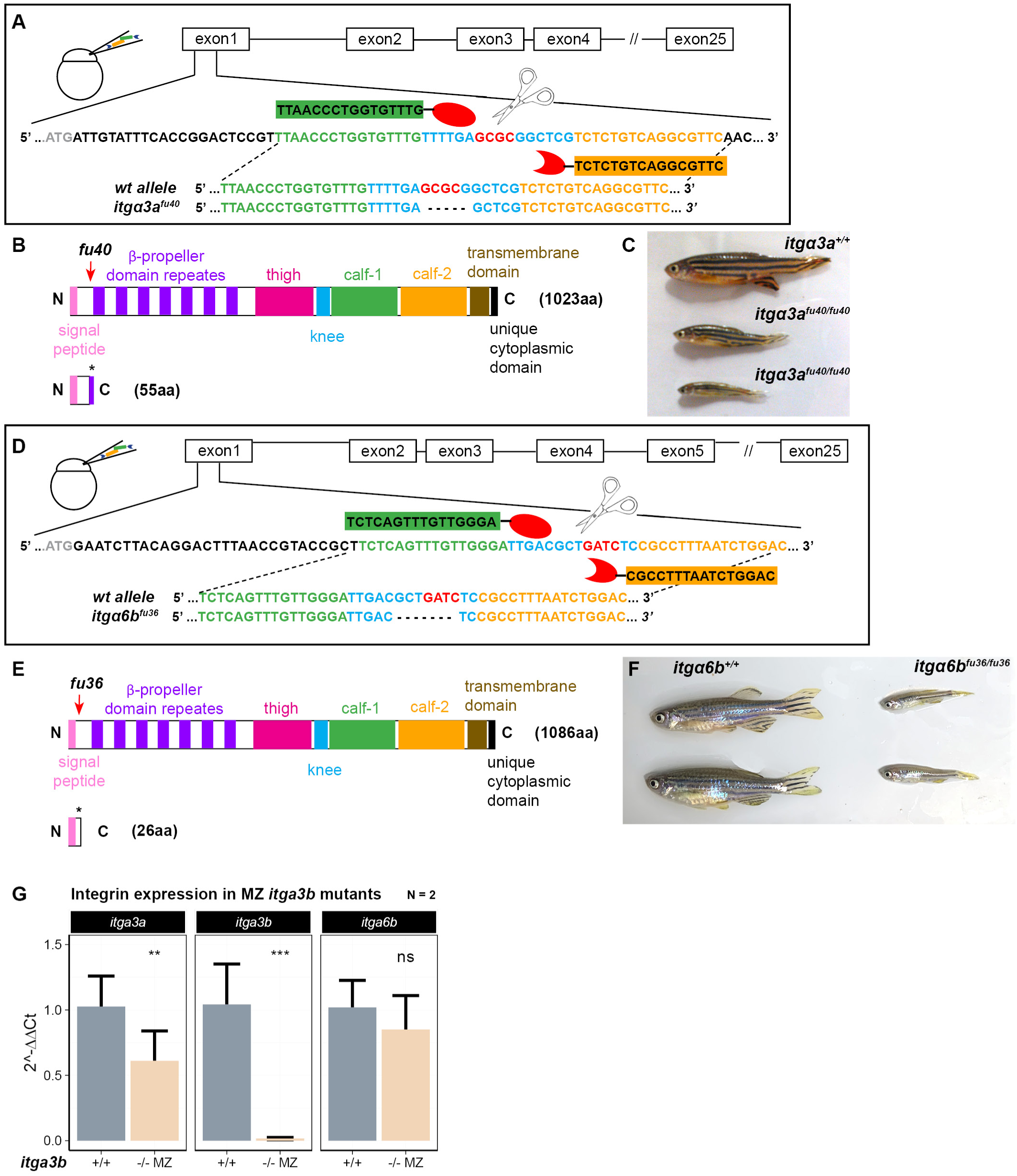
Integrin mutant generation and qPCR analysis. (A-B, D-E) Schematic of *itga3a* and *itga6b* TALEN design and predicted truncated proteins. (C, F) Rare escapers of *itga3a* and *itga6b* homozygous mutants with severe dwarfism. (G) qRT-PCR expression analysis of *itga3a, itga3b* and *itga6b* in maternal zygotic (MZ) *itga3b* mutants.

**S3:**
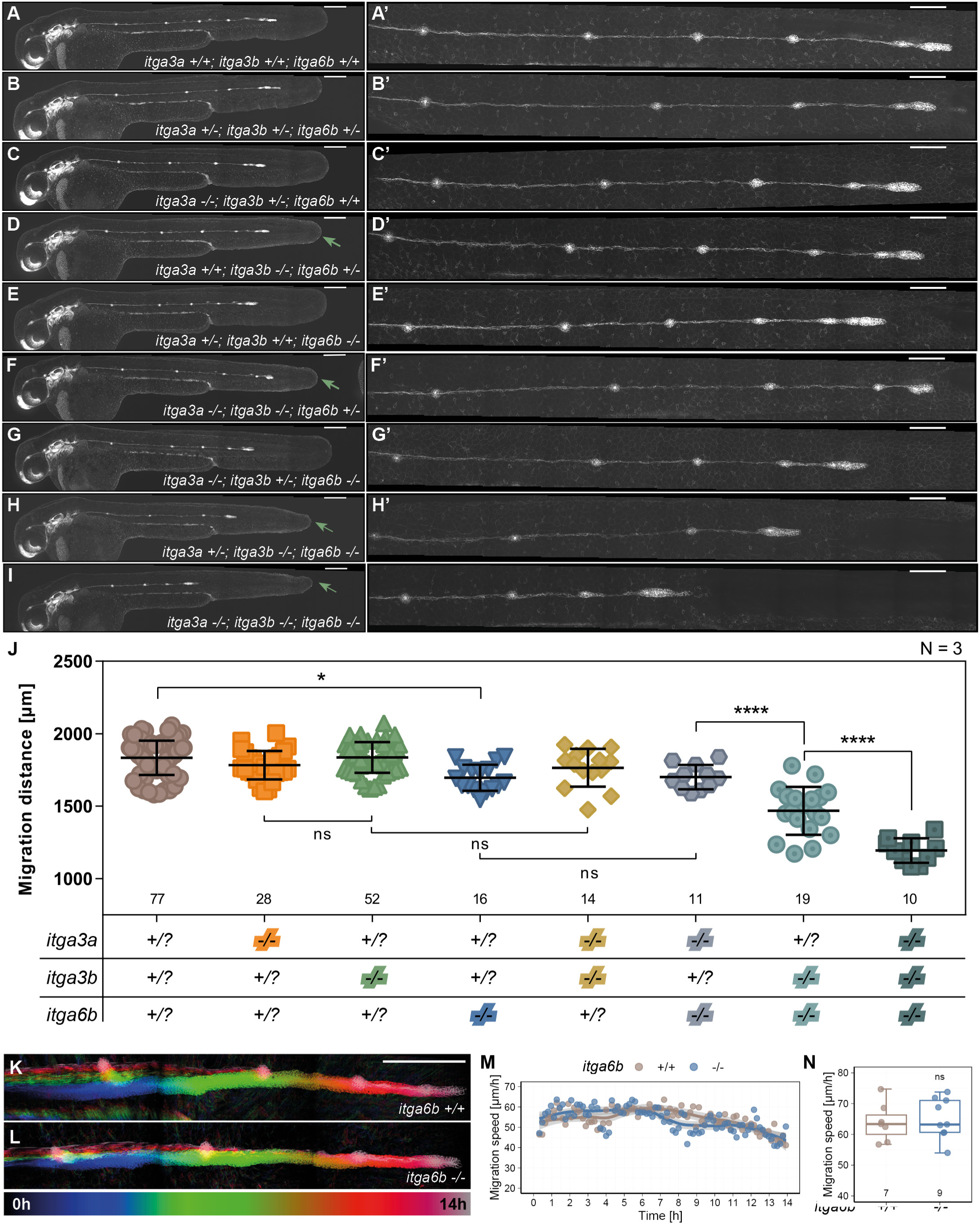
Integrin mutant analysis. (A-I’) Full panel of triple integrin mutant combinations from Figure 3, including embryo overviews (A-I) and higher magnifications of the pLL (A’-I’). (J) Quantification of the distance migrated by the pLLP at 40 hpf. (K-L) Temporally color-coded maximum intensity projection (MIPs) from 14-hour TLs of wild-type and *itga6b* mutant embryos. (M-N) Quantification of instantaneous and mean speeds. Statistics: One-way ANOVA with multiple comparisons (J) and two-tailed Wilcoxon rank-sum test (N); ns = non-significant; *p < 0.05; ****p < 0.0001. Sample sizes (n) are indicated above x-axes. N = number of biological replicates. Scale bars: 200µm (A-I, K-L), 100µm (A’-I’).

**S4:**
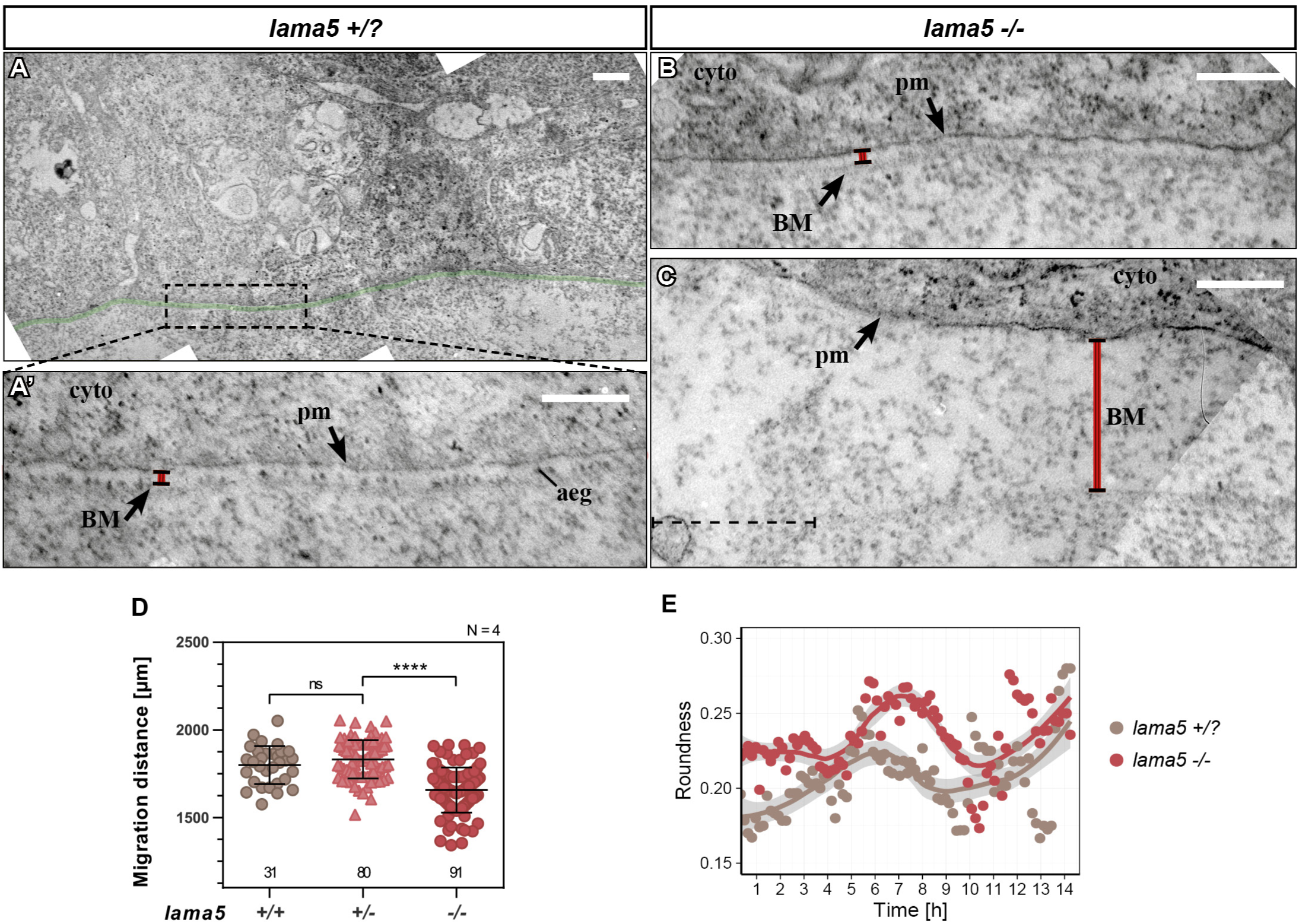
*lama5* mutant analysis. (A-C) TEM in 32hpf sibling (A) and *lama5^−/−^* (B,C) embryos. (A) Shown is the pLLP leading region with the underlying epidermal BM (green). (A’) Close-up as indicated in (A) showing the BM spanning ∼50nm and containing first adepidermal granules (aeg). (B) In the BM of *lama5* mutants, adepidermal granules are missing from intact BM regions, where the width is comparable to control siblings (A). (C) In the proximity of BM holes (dashed line), the BM is bloated. (D) Quantification of the migrated distance at 40hpf including heterozygous individuals. (E) Quantification of roundness over time in control and *lama5* mutants. Statistics: One-way ANOVA with multiple comparisons; ns = non-significant; ****p < 0.0001. Scale bars: 500nm (A) and 250nm (A’, B, C). cyto = cytoplasm, pm = plasma membrane, BM = basement membrane, aeg = adepidermal granule.

**S5:**
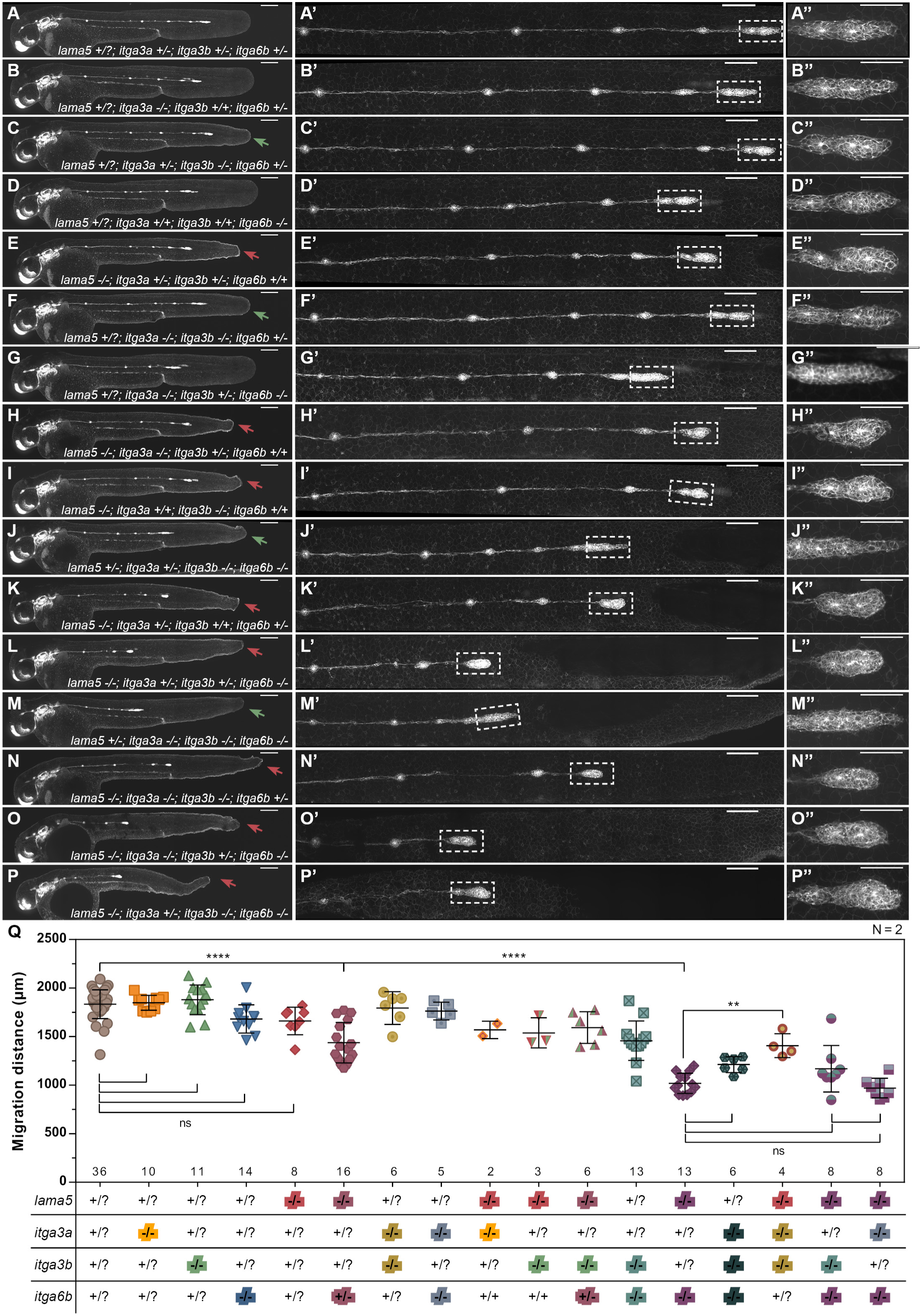
Analysis of laminin-integrin mutant combinations. (A-P’’) Full panel of *lama5*-integrin mutant combinations from Figure 7, including embryo overviews (A-P), higher magnifications of the pLL (A’-P’) and close-ups on the pLLP (A’’-P’’). (Q) Quantification of the distance migrated by the pLLP at 40 hpf. Statistics: One-way ANOVA with multiple comparisons; ns = non-significant; *p < 0.05; ****p < 0.0001. Sample sizes (n) are indicated above x-axes. N = number of biological replicates. Scale bars: 200µm (A-P), 100µm (A’-P’) and 50µm (A’’-P’’).

**S6:**
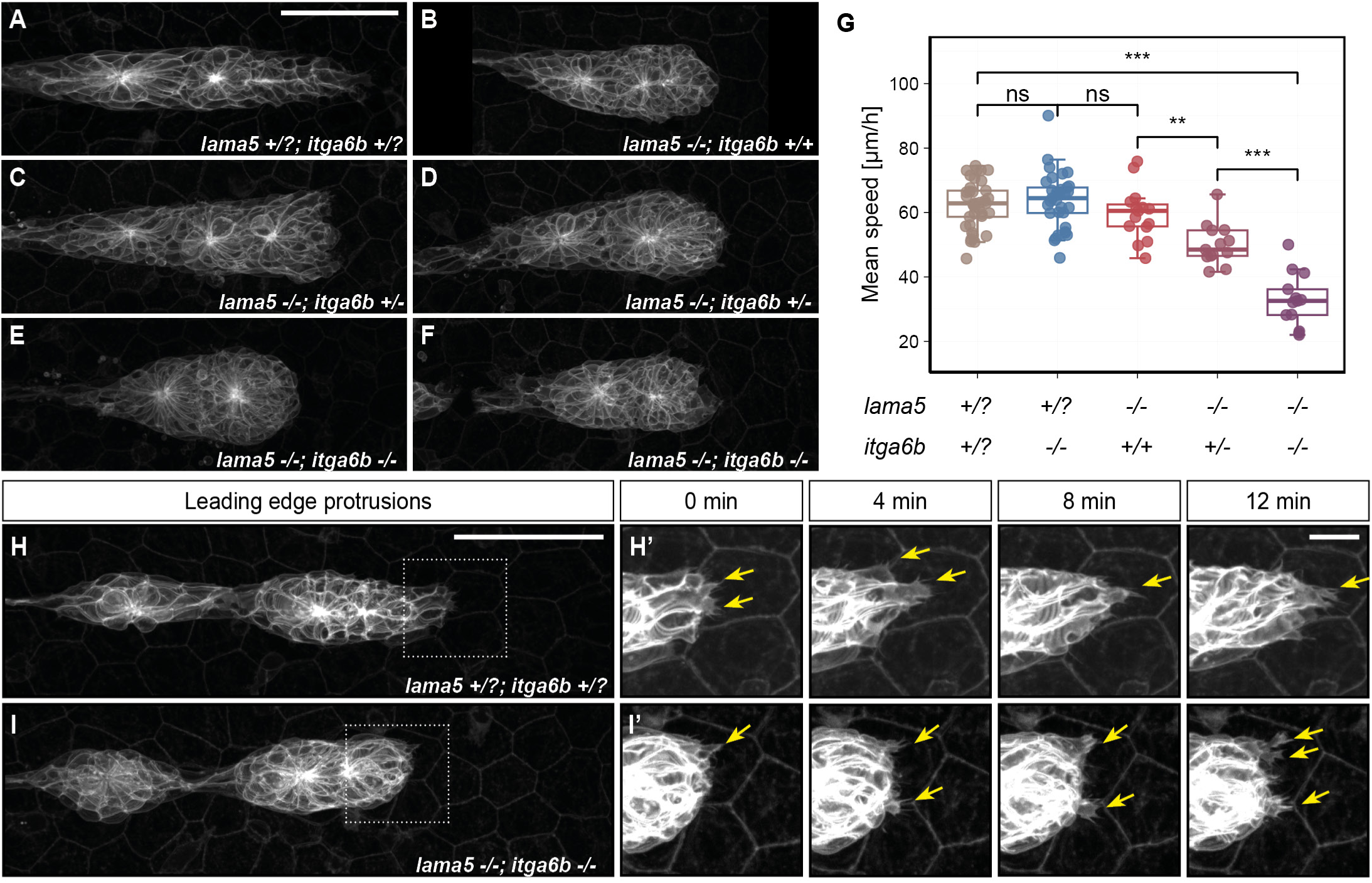
*lama5;itga6b* mutants. (A-F) Live imaging of pLLPs of representative sibling control (A), *lama5* single mutant (B), *lama5^−/−^;itga6b^+/-^* (C-D) and *lama5^−/−^;itga6b^−/−^* double mutants (E-F). (G) Full quantification of mean speeds shown in Figure 7L (H-J’) Time-lapse imaging of dynamic leading edge protrusions in control (H-H’) and *lama5;itga6b* double mutant pLLPs (J-J’). Scale bars: 50µm (A, H), 10µm (H’).

## Materials and Methods

### Zebrafish

Procedures involving animals were carried out according to the guidelines of the Goethe University of Frankfurt and were approved by the German authorities (Veterinary Department of the Regional Board of Darmstadt). Zebrafish embryos were obtained through natural spawning and staged as previously described ^37^. *bdf^fr^*^21^ (*badfin/itga3b*) and *fra^tc^*^17^ (*fransen/lama5*) ^39^ mutant lines and *Tg(-0.8cldnb:lynEGFP)^zf^*^106^ (referred to as *cldnb:GFP*, Haas and Gilmour, 2006), *TgBAC(lamc1:lamc1-sfGFP)^sk^*^116^*^Tg^* (referred to as *lamc1-GFP*, Yamaguchi *et al.*, 2022), *Tg(cxcr4b(FOS):Lifeact-mRuby)^fu^*^16^ (referred to as *cxcr4b:R-Lifeact*, Lardennois *et al.*, 2025, bioRxiv) transgenic lines were described previously. The alleles *itga3a^fu^*^40^ and *itga6b^fu^*^36^ were generated using TALEN targeting exon 1 as previously described ^76,77^.

### Genotyping

To genotype mutant alleles, gDNA was extracted from fins or single embryos by boiling at 95°C in 50 mM NaOH and neutralizing with 1/10 volume 1 M Tris-HCl (pH 8.0). Allele-specific PCRs were carried out using 1 µL gDNA using the primers and according annealing temperatures listed in Table S1. Following PCR, the *fra^tc^*^17^ allele was genotyped by sequencing to identify the single nucleotide change (AAAAA > AATAA). Sequencing was performed by Eurofins GATC Supreme Run Sequencing. The *itga3b^fr^*^21^ allele was genotyped with the dCAPS (derived Cleaved Amplified Polymorphic Sequences) technique ^78^ and digest with EcoRV-HF yielding 231bp/26bp products. The *itga3a^fu^*^40^ allele was genotyped by digest with HhaI yielding 240bp/198bp products. The *itga6b^fu^*^36^ allele was genotyped by digest with MboI yielding 353bp/212bp products.

### RNA extraction, cDNA synthesis and qPCR

RNA extraction was performed using TRIzol reagent according to manufacturer’s instructions. Embryos at 32hpf were dechorionated and homogenized in 500μL TRIzol with a blunt needle tip syringe (20-40 embryos per tube).

cDNA was generated using 1 µg of total RNA with the Invitrogen SuperscriptIII® First-Strand Synthesis System for ISH probe synthesis and iScriptTM cDNA Synthesis Kit (Bio-Rad, 1708890) for qPCR, according to manufacturer’s instructions.

For qPCR, cDNA was diluted 1:1. Primers for qPCR were designed using the Primer3-based Primer-BLAST tool from the NCBI. Real-time qPCR was performed using the Bio-Rad iTaq Universal SYBR Green Supermix (1725270) according to manufacturers instructions in a CFX Connect Real-Time System (Biorad) running 40 cycles of 15 sec denaturation (95°C) and 60 sec annealing+extension (60°C). Fold changes were calculated using the 2^ΔΔCt^ method ^79^. Expression of the genes of interest were normalized to that of *rpl13*.

### Embryo fixation, *in-situ* hybridization and immunostainings

In general, embryos were fixed overnight at 4°C or 2h at room temperature with 4% PFA in 1x PBS. After fixation, embryos were washed in PBS and either used directly for experiments or dehydrated in methanol/PBS series and stored at -20°C for *in-situ* hybridization (ISH).

Wholemount immunostaining was performed according to standard procedures ^80^. The following primary antibodies were used: anti-GFP (mouse monoclonal, 1:500, Takara Bio Cat# 632380, RRID:AB_10013427), anti-pan-laminin (rabbit, 1:400, Sigma-Aldrich Cat# L9393, RRID:AB_477163), anti-pFAK1 (rabbit, 1:200, Invitrogen, Cat# 44-624G), anti-lama5 (rabbit, 1:200, #504, Hannocks *et al.*, 2018). Anti-rabbit and anti-mouse Alexa conjugated antibodies were used as secondary antibodies (1:500, Molecular Probes, Eugene, OR, USA).

To generate riboprobes for ISH, partial cDNA sequences for *itga3a*, *itga3b*, *itga6a*, *itga6b*, *itga6l*, *itgb1a* and *itgb1b* were amplified by PCR using the primers listed in Table S1. PCR products were subcloned in pGEM-T-easy (Promega) and Digoxigenin(DIG)-labelled riboprobes were transcribed from linearized plasmid using Sp6 (Thermo Fisher) or T7 polymerases (Thermo Fisher). ISH was performed according to standard procedures ^80^ with probes diluted 1:100 and followed by immunostaining using anti-GFP antibody as explained above.

### Imaging

#### Mounting

For live imaging, embryos were dechorionated using pronase and mounted in 0.3% low-melting agarose (LMPA) in an agarose cast ^81^, with the following modifications: no second layer of 0.5% LMPA was added after polymerization.

#### ISH images

Imaging of ISH embryos was performed on an inverted microscope (Ti-S, Nikon) using a 4x air objective to acquire whole embryos and a 20x air objective for close-ups of the pLLP.

#### Spinning disc

Imaging of live and fixed embryos was performed on a Nikon W1 Spinning Disc Microscope with the following objectives: 10× air objective (NA 0.45, WD 4 mm), 20× (NA 0.95,WD 0.95 mm), and 40× (NA 1.15,WD 0.60 mm) water objectives.

For 14-h timelapse recordings, embryos were selected at 26hpf-30hpf, mounted as described above and imaged using the 20x objective with additional 1.5x digital zoom. To ensure water-immersion of the objective over the course of the timelapse, a water-dispenser was manually installed. An incubation chamber surrounding the microscope kept embryos at 28°C during the experiment. The 488 nm laser was set to 20% power, 30 ms exposure time, 2×2 binning and gain 4. Images were acquired as z-stacks with 5 µm z-spacing at 10 min frame interval.

### Image processing

Images were processed using the open-source processing software Fiji (Fiji Is Just ImageJ) (2.1.0/1.53c). For presentation purposes, three-dimensional z-stacks were generally flattened by maximum intensity projection (ZPMAX). For immunostainings, single z-slices were selected, and signal intensities were adjusted for better visualization of the antibody signal. In experiments with control and mutant groups, signal intensities were adjusted in the same way for all images. Quantifications of intensity, length or roundness were performed using segmented line or freehand selection tools in Fiji. Quantifications performed in Fiji were further processed and plotted in R (version 4.3.1) using the ggplot2 package (version 3.4.3) or GraphPad Prism (version 6.07).

### Segmentation and analysis of 14h TLs

Analysis of 14-h timelapses was performed using a two-part custom-written macro in Fiji. First, the migrating pLLP and deposited neuromasts were automatically segmented from each frame using “MaxEntropy” thresholding, generating a binary video. Missegmentations due to the skin and kidney also being labelled by *cldnb:GFP*, were manually corrected by the user. Second, this binary movie was subjected to the “Analyze particles” plug-in and the pLLP was registered as the particle with highest X coordinate. Finally, the position of the pLLP leading edge was determined using bounding rectangle coordinates and roundness was measured. The instantaneous speed between frames and mean speed was then calculated in R. Plots of speed and roundness over-time were generated using the ggplot2 package and trends visualized by locally estimated scatterplot smoothing (loess) curves. Mean speeds were plotted as boxplots overlayed by individual datapoints.

### Statistical analysis

Statistical analysis in R was performed using the ggpubr package (version 0.6.0) with `‘stat_compare_means’` function or `‘geom_signif’` function for pairwise comparisons. Unless otherwise stated, the non-parametric two-sided Wilcoxon rank-sum test (also known as Mann-Whitney U test) was used. For pairwise comparisons, *p*-values were adjusted for multiple testing using the Holm correction method.

In GraphPad Prism, normality was assessed using the Shapiro-Wilk test, and if confirmed, comparisons were made using one-way ANOVA with or without multiple comparisons. If normality was not confirmed, the Mann-Whitney U test was used.

Statistical significance was defined as *p* < 0.05. ns = not significant; *\*p* < 0.05, *\*\*p* < 0.01, and *\*\*\*p* < 0.001, *\*\*\*\*p* < 0.0001.

## Bibliography

1. Lecaudey, V., and Gilmour, D. (2006). Organizing moving groups during morphogenesis. Current Opinion in Cell Biology 18, 102–107. 10.1016/j.ceb.2005.12.001.

2. Friedl, P., and Gilmour, D. (2009). Collective cell migration in morphogenesis, regeneration and cancer. Nat Rev Mol Cell Biol 10, 445–457. 10.1038/nrm2720.

3. Norden, C., and Lecaudey, V. (2019). Collective cell migration: general themes and new paradigms. Current Opinion in Genetics & Development 57, 54–60. 10.1016/j.gde.2019.06.013.

4. De Pascalis, C., and Etienne-Manneville, S. (2017). Single and collective cell migration: the mechanics of adhesions. MBoC 28, 1833–1846. 10.1091/mbc.e17-03-0134.

5. Chastney, M.R., Kaivola, J., Leppänen, V.-M., and Ivaska, J. (2024). The role and regulation of integrins in cell migration and invasion. Nat Rev Mol Cell Biol. 10.1038/s41580-024-00777-1.

6. Hynes, R.O. (2002). Integrins. Cell 110, 673–687. 10.1016/S0092-8674(02)00971-6.

7. Takada, Y., Ye, X., and Simon, S. (2007). The integrins. Genome Biol 8, 215. 10.1186/gb-2007-8-5-215.

8. Bachmann, M., Kukkurainen, S., Hytönen, V.P., and Wehrle-Haller, B. (2019). Cell Adhesion by Integrins. Physiological Reviews 99, 1655–1699. 10.1152/physrev.00036.2018.

9. Yamada, K.M., and Sixt, M. (2019). Mechanisms of 3D cell migration. Nat Rev Mol Cell Biol 20, 738–752. 10.1038/s41580-019-0172-9.

10. Scarpa, E., and Mayor, R. (2016). Collective cell migration in development. Journal of Cell Biology 212, 143–155. 10.1083/jcb.201508047.

11. Alberts, B. (2015). Molecular biology of the cell Sixth edition. (Garland Science, Taylor and Francis Group).

12. Sherwood, D.R. (2021). Basement membrane remodeling guides cell migration and cell morphogenesis during development. Current Opinion in Cell Biology 72, 19–27. 10.1016/j.ceb.2021.04.003.

13. Walma, D.A.C., and Yamada, K.M. (2020). The extracellular matrix in development. Development 147, dev175596. 10.1242/dev.175596.

14. Hohenester, E., and Yurchenco, P.D. (2013). Laminins in basement membrane assembly. Cell Adh Migr 7, 56–63. 10.4161/cam.21831.

15. Belkin, A.M., and Stepp, M.A. (2000). Integrins as receptors for laminins. Microsc. Res. Tech. 51, 280–301. 10.1002/1097-0029(20001101)51:3%253C280::AID-JEMT7%253E3.0.CO;2-O.

16. Klaffky, E., Williams, R., Yao, C.-C., Ziober, B., Kramer, R., and Sutherland, A. (2001). Trophoblast-Specific Expression and Function of the Integrin α7 Subunit in the Peri-implantation Mouse Embryo. Developmental Biology 239, 161–175. 10.1006/dbio.2001.0404.

17. Li, S., Edgar, D., Fässler, R., Wadsworth, W., and Yurchenco, P.D. (2003). The Role of Laminin in Embryonic Cell Polarization and Tissue Organization. Developmental Cell 4, 613–624. 10.1016/S1534-5807(03)00128-X.

18. Miner, J.H., Cunningham, J., and Sanes, J.R. (1998). Roles for laminin in embryogenesis: exencephaly, syndactyly, and placentopathy in mice lacking the laminin alpha5 chain. J Cell Biol 143, 1713–1723. 10.1083/jcb.143.6.1713.

19. Miner, J.H. (2004). Compositional and structural requirements for laminin and basement membranes during mouse embryo implantation and gastrulation. Development 131, 2247–2256. 10.1242/dev.01112.

20. Spenlé, C., Simon-Assmann, P., Orend, G., and Miner, J.H. (2013). Laminin α5 guides tissue patterning and organogenesis. Cell Adhesion & Migration 7, 90–100. 10.4161/cam.22236.

21. Gordon-Weeks, A., Lim, S.Y., Yuzhalin, A., Lucotti, S., Vermeer, J.A.F., Jones, K., Chen, J., and Muschel, R.J. (2019). Tumour-Derived Laminin α5 (LAMA5) Promotes Colorectal Liver Metastasis Growth, Branching Angiogenesis and Notch Pathway Inhibition. Cancers 11, 630. 10.3390/cancers11050630.

22. Kikkawa, Y., Ogawa, T., Sudo, R., Yamada, Y., Katagiri, F., Hozumi, K., Nomizu, M., and Miner, J.H. (2013). The Lutheran/Basal Cell Adhesion Molecule Promotes Tumor Cell Migration by Modulating Integrin-mediated Cell Attachment to Laminin-511 Protein. Journal of Biological Chemistry 288, 30990–31001. 10.1074/jbc.m113.486456.

23. Qin, Y., Shembrey, C., Smith, J., Paquet-Fifield, S., Behrenbruch, C., Beyit, L.M., Thomson, B.N.J., Heriot, A.G., Cao, Y., and Hollande, F. (2020). Laminin 521 enhances self-renewal via STAT3 activation and promotes tumor progression in colorectal cancer. Cancer Letters 476, 161–169. 10.1016/j.canlet.2020.02.026.

24. Barraza-Flores, P., Bates, C.R., Oliveira-Santos, A., and Burkin, D.J. (2020). Laminin and Integrin in LAMA2-Related Congenital Muscular Dystrophy: From Disease to Therapeutics. Front. Mol. Neurosci. 13, 1. 10.3389/fnmol.2020.00001.

25. Has, C., Spartà, G., Kiritsi, D., Weibel, L., Moeller, A., Vega-Warner, V., Waters, A., He, Y., Anikster, Y., Esser, P., et al. (2012). Integrin α_3_ Mutations with Kidney, Lung, and Skin Disease. N Engl J Med 366, 1508–1514. 10.1056/NEJMoa1110813.

26. Ridley, A.J., Schwartz, M.A., Burridge, K., Firtel, R.A., Ginsberg, M.H., Borisy, G., Parsons, J.T., and Horwitz, A.R. (2003). Cell Migration: Integrating Signals from Front to Back. Science 302, 1704–1709. 10.1126/science.1092053.

27. Vaughan, R.B., and Trinkaus, J.P. (1966). Movements of epithelial cell sheets *in vitro*. Journal of Cell Science 1, 407–413. 10.1242/jcs.1.4.407.

28. Weiner, O.D., Servant, G., Welch, M.D., Mitchison, T.J., Sedat, J.W., and Bourne, H.R. (1999). Spatial control of actin polymerization during neutrophil chemotaxis. Nat Cell Biol 1, 75–81. 10.1038/10042.

29. Mayor, R., and Etienne-Manneville, S. (2016). The front and rear of collective cell migration. Nat Rev Mol Cell Biol 17, 97–109. 10.1038/nrm.2015.14.

30. Sun, Z., Guo, S.S., and Fässler, R. (2016). Integrin-mediated mechanotransduction. Journal of Cell Biology 215, 445–456. 10.1083/jcb.201609037.

31. Ladoux, B., and Mège, R.-M. (2017). Mechanobiology of collective cell behaviours. Nat Rev Mol Cell Biol 18, 743–757. 10.1038/nrm.2017.98.

32. Dalle Nogare, D., and Chitnis, A.B. (2017). A framework for understanding morphogenesis and migration of the zebrafish posterior Lateral Line primordium. Mechanisms of Development 148, 69–78. 10.1016/j.mod.2017.04.005.

33. Ghysen, A., and Dambly-Chaudiere, C. (2007). The lateral line microcosmos. Genes & Development 21, 2118–2130. 10.1101/gad.1568407.

34. Durdu, S., Iskar, M., Revenu, C., Schieber, N., Kunze, A., Bork, P., Schwab, Y., and Gilmour, D. (2014). Luminal signalling links cell communication to tissue architecture during organogenesis. Nature 515, 120–124. 10.1038/nature13852.

35. Lecaudey, V., Cakan-Akdogan, G., Norton, W.H.J., and Gilmour, D. (2008). Dynamic Fgf signaling couples morphogenesis and migration in the zebrafish lateral line primordium. Development 135, 2695–2705. 10.1242/dev.025981.

36. Yamaguchi, N., Zhang, Z., Schneider, T., Wang, B., Panozzo, D., and Knaut, H. (2022). Rear traction forces drive adherent tissue migration in vivo. Nat Cell Biol 24, 194–204. 10.1038/s41556-022-00844-9.

37. Kimmel, C.B., Ballard, W.W., Kimmel, S.R., Ullmann, B., and Schilling, T.F. (1995). Stages of embryonic development of the zebrafish. Dev Dyn 203, 253–310. 10.1002/aja.1002030302.

38. Haas, P., and Gilmour, D. (2006). Chemokine Signaling Mediates Self-Organizing Tissue Migration in the Zebrafish Lateral Line. Developmental Cell 10, 673–680. 10.1016/j.devcel.2006.02.019.

39. Carney, T.J., Kiyozumi, D., Gebauer, J.M., Talbot, J.C., Kimmel, C.B., Wagener, R., Schwarz, H., Ingham, P.W., and Hammerschmidt, M. (2010). Genetic Analysis of Fin Development in Zebrafish Identifies Furin and Hemicentin1 as Potential Novel Fraser Syndrome Disease Genes. PLoS Genetics 6, 22.

40. El-Brolosy, M.A., Kontarakis, Z., Rossi, A., Kuenne, C., Günther, S., Fukuda, N., Kikhi, K., Boezio, G.L.M., Takacs, C.M., Lai, S.-L., et al. (2019). Genetic compensation triggered by mutant mRNA degradation. Nature 568, 193–197. 10.1038/s41586-019-1064-z.

41. Jülich, D., Cobb, G., Melo, A.M., McMillen, P., Lawton, A.K., Mochrie, S.G.J., Rhoades, E., and Holley, S.A. (2015). Cross-Scale Integrin Regulation Organizes ECM and Tissue Topology. Developmental Cell 34, 33–44. 10.1016/j.devcel.2015.05.005.

42. Schober, M., Raghavan, S., Nikolova, M., Polak, L., Pasolli, H.A., Beggs, H.E., Reichardt, L.F., and Fuchs, E. (2007). Focal adhesion kinase modulates tension signaling to control actin and focal adhesion dynamics. The Journal of Cell Biology 176, 667–680. 10.1083/jcb.200608010.

43. Nishiuchi, R., Takagi, J., Hayashi, M., Ido, H., Yagi, Y., Sanzen, N., Tsuji, T., Yamada, M., and Sekiguchi, K. (2006). Ligand-binding specificities of laminin-binding integrins: A comprehensive survey of laminin–integrin interactions using recombinant α3β1, α6β1, α7β1 and α6β4 integrins. Matrix Biology 25, 189–197. 10.1016/j.matbio.2005.12.001.

44. Arimori, T., Miyazaki, N., Mihara, E., Takizawa, M., Taniguchi, Y., Cabañas, C., Sekiguchi, K., and Takagi, J. (2021). Structural mechanism of laminin recognition by integrin. Nat Commun 12, 4012. 10.1038/s41467-021-24184-8.

45. Hannocks, M.-J., Pizzo, M.E., Huppert, J., Deshpande, T., Abbott, N.J., Thorne, R.G., and Sorokin, L. (2018). Molecular characterization of perivascular drainage pathways in the murine brain. J Cereb Blood Flow Metab 38, 669–686. 10.1177/0271678X17749689.

46. Bader, H.L., Lambert, E., Guiraud, A., Malbouyres, M., Driever, W., Koch, M., and Ruggiero, F. (2013). Zebrafish Collagen XIV Is Transiently Expressed in Epithelia and Is Required for Proper Function of Certain Basement Membranes. Journal of Biological Chemistry 288, 6777–6787. 10.1074/jbc.M112.430637.

47. Ghysen, A., and Dambly-Chaudière, C. (2004). Development of the zebrafish lateral line. Current Opinion in Neurobiology 14, 67–73. 10.1016/j.conb.2004.01.012.

48. Yurchenco, P.D., Quan, Y., Colognato, H., Mathus, T., Harrison, D., Yamada, Y., and O’Rear, J.J. (1997). The α chain of laminin-1 is independently secreted and drives secretion of its β- and γ-chain partners. Proc. Natl. Acad. Sci. U.S.A. 94, 10189–10194. 10.1073/pnas.94.19.10189.

49. Aumailley, M. (2013). The laminin family. Cell Adhesion & Migration 7, 48–55. 10.4161/cam.22826.

50. Parsons, M.J., Pollard, S.M., Saúde, L., Feldman, B., Coutinho, P., Hirst, E.M.A., and Stemple, D.L. (2002). Zebrafish mutants identify an essential role for laminins in notochord formation. Development (Cambridge, England) 129, 3137—3146.

51. Sztal, T., Berger, S., Currie, P.D., and Hall, T.E. (2011). Characterization of the laminin gene family and evolution in zebrafish. Dev. Dyn. 240, 422–431. 10.1002/dvdy.22537.

52. Farnsworth, D.R., Saunders, L.M., and Miller, A.C. (2020). A single-cell transcriptome atlas for zebrafish development. Developmental Biology 459, 100–108. 10.1016/j.ydbio.2019.11.008.

53. Sur, A., Wang, Y., Capar, P., Margolin, G., Prochaska, M.K., and Farrell, J.A. (2023). Single-cell analysis of shared signatures and transcriptional diversity during zebrafish development. Developmental Cell 58, 3028–3047.e12. 10.1016/j.devcel.2023.11.001.

54. Fukumoto, S., Miner, J.H., Ida, H., Fukumoto, E., Yuasa, K., Miyazaki, H., Hoffman, M.P., and Yamada, Y. (2006). Laminin α5 Is Required for Dental Epithelium Growth and Polarity and the Development of Tooth Bud and Shape. Journal of Biological Chemistry 281, 5008–5016. 10.1074/jbc.M509295200.

55. Webb, A.E., Sanderford, J., Frank, D., Talbot, W.S., Driever, W., and Kimelman, D. (2007). Laminin α5 is essential for the formation of the zebrafish fins. Developmental Biology 311, 369–382. 10.1016/j.ydbio.2007.08.034.

56. Miner, J.H., Li, C., Mudd, J.L., Go, G., and Sutherland, A.E. (2004). Compositional and structural requirements for laminin and basement membranes during mouse embryo implantation and gastrulation. Development 131, 2247–2256. 10.1242/dev.01112.

57. Mikdache, A., Boueid, M.-J., Lesport, E., Delespierre, B., Loisel-Duwattez, J., Degerny, C., and Tawk, M. (2022). Timely Schwann cell division drives peripheral myelination *in vivo* via the laminin/cAMP pathway. Development 149, dev200640. 10.1242/dev.200640.

58. Meighan, C.M., and Schwarzbauer, J.E. (2008). Temporal and spatial regulation of integrins during development. Current Opinion in Cell Biology 20, 520–524. 10.1016/j.ceb.2008.05.010.

59. Truong, H., and Danen, E.H.J. (2009). Integrin switching modulates adhesion dynamics and cell migration. Cell Adhesion & Migration 3, 179–181. 10.4161/cam.3.2.8036.

60. Madamanchi, A., Zijlstra, A., and Zutter, M.M. (2014). Flipping the Switch: Integrin Switching Provides Metastatic Competence. Sci. Signal. 7. 10.1126/scisignal.2005236.

61. Winkler, J., Abisoye-Ogunniyan, A., Metcalf, K.J., and Werb, Z. (2020). Concepts of extracellular matrix remodelling in tumour progression and metastasis. Nat Commun 11, 5120. 10.1038/s41467-020-18794-x.

62. Bair, E.L., Chen, M.L., McDaniel, K., Sekiguchi, K., Cress, A.E., Nagle, R.B., and Bowden, G.T. (2005). Membrane Type 1 Matrix Metalloprotease Cleaves Laminin-10 and Promotes Prostate Cancer Cell Migration. Neoplasia 7, 380–389. 10.1593/neo.04619.

63. Gopal, S.K., Greening, D.W., Zhu, H.-J., Simpson, R.J., and Mathias, R.A. (2016). Transformed MDCK cells secrete elevated MMP1 that generates LAMA5 fragments promoting endothelial cell angiogenesis. Sci Rep 6, 28321. 10.1038/srep28321.

64. Niland, S., and Eble, J.A. (2025). Decoding the MMP14 integrin link: Key player in the secretome landscape. Matrix Biology 136, 36–51. 10.1016/j.matbio.2025.01.004.

65. Burkin, D.J., and Kaufman, S.J. (1999). The α7β1 integrin in muscle development and disease. Cell and Tissue Research 296, 183–190. 10.1007/s004410051279.

66. Yeh, M.-G., Ziober, B.L., Liu, B., Lipkina, G., Vizirianakis, I.S., and Kramer, R.H. (2003). The beta1 cytoplasmic domain regulates the laminin-binding specificity of the alpha7X1 integrin. Mol Biol Cell 14, 3507–3518. 10.1091/mbc.e02-12-0824.

67. Brooks, D.L.P., Schwab, L.P., Krutilina, R., Parke, D.N., Sethuraman, A., Hoogewijs, D., Schörg, A., Gotwald, L., Fan, M., Wenger, R.H., et al. (2016). ITGA6 is directly regulated by hypoxia-inducible factors and enriches for cancer stem cell activity and invasion in metastatic breast cancer models. Mol Cancer 15, 26. 10.1186/s12943-016-0510-x.

68. Kwon, J., Lee, T.-S., Lee, H.W., Kang, M.C., Yoon, H.-J., Kim, J.-H., and Park, J.H. (2013). Integrin alpha 6: A novel therapeutic target in esophageal squamous cell carcinoma. International Journal of Oncology 43, 1523–1530. 10.3892/ijo.2013.2097.

69. Ma, G., Jing, C., Huang, F., Li, X., Cao, X., and Liu, Z. (2017). Integrin α6 promotes esophageal cancer metastasis and is targeted by miR-92b. Oncotarget 8, 6681–6690. 10.18632/oncotarget.14259.

70. Zheng, G., Bouamar, H., Cserhati, M., Zeballos, C.R., Mehta, I., Zare, H., Broome, L., Hu, R., Lai, Z., Chen, Y., et al. (2022). Integrin alpha 6 is upregulated and drives hepatocellular carcinoma progression through integrin α6β4 complex. Intl Journal of Cancer 151, 930–943. 10.1002/ijc.34146.

71. Shellard, A., Szabó, A., Trepat, X., and Mayor, R. (2018). Supracellular contraction at the rear of neural crest cell groups drives collective chemotaxis. Science 362, 339–343. 10.1126/science.aau3301.

72. Valentin, G., Haas, P., and Gilmour, D. (2007). The Chemokine SDF1a Coordinates Tissue Migration through the Spatially Restricted Activation of Cxcr7 and Cxcr4b. Current Biology 17, 1026–1031. 10.1016/j.cub.2007.05.020.

73. Nechiporuk, A., and Raible, D.W. (2008). FGF-Dependent Mechanosensory Organ Patterning in Zebrafish. Science 320, 1774–1777. 10.1126/science.1156547.

74. Mukherjee, A., Hilzendeger, M., Rinvelt, A., Fatma, S., Schupp, M., Dalle Nogare, D., and Chitnis, A.B. (2025). Signaling and Mechanics influence the number and size of epithelial rosettes in the migrating zebrafish Posterior Lateral Line primordium. Preprint at Developmental Biology, 10.1101/2025.05.17.650326 10.1101/2025.05.17.650326.

75. Lardennois, A., Dingare, C., Duda, V., Klemmt, P.A., Heinzen, C., Kleinhans, D., Falk, T., Schelmbauer, C., Mozolewska, O., Papadopoulou, S., et al. (2025). Vgll4 Proteins limit Organ Size in Zebrafish through Yap1-Dependent and -Independent Mechanisms. Preprint at Developmental Biology, 10.1101/2025.05.08.652796 10.1101/2025.05.08.652796.

76. Agarwala, S., Duquesne, S., Liu, K., Boehm, A., Grimm, L., Link, S., König, S., Eimer, S., Ronneberger, O., and Lecaudey, V. (2015). Amotl2a interacts with the Hippo effector Yap1 and the Wnt/β-catenin effector Lef1 to control tissue size in zebrafish. eLife 4, e08201. 10.7554/eLife.08201.

77. Dingare, C., Niedzwetzki, A., Klemmt, P.A., Godbersen, S., Fuentes, R., Mullins, M.C., and Lecaudey, V. (2018). The Hippo pathway effector Taz is required for cell morphogenesis and fertilization in zebrafish. Development, dev.167023. 10.1242/dev.167023.

78. Neff, M.M., Neff, J.D., Chory, J., and Pepper, A.E. (1998). dCAPS, a simple technique for the genetic analysis of single nucleotide polymorphisms: experimental applications in *Arabidopsis thaliana* genetics. The Plant Journal 14, 387–392. 10.1046/j.1365-313X.1998.00124.x.

79. Livak, K.J., and Schmittgen, T.D. (2001). Analysis of Relative Gene Expression Data Using Real-Time Quantitative PCR and the 2−ΔΔCT Method. Methods 25, 402–408. 10.1006/meth.2001.1262.

80. Lecaudey, V., Anselme, I., Rosa, F., and Schneider-Maunoury, S. (2004). The zebrafish Iroquois gene *iro7* positions the r4/r5 boundary and controls neurogenesis in the rostral hindbrain. Development 131, 3121–3131. 10.1242/dev.01190.

81. Kleinhans, D.S., and Lecaudey, V. (2019). Standardized mounting method of (zebrafish) embryos using a 3D-printed stamp for high-content, semi-automated confocal imaging. BMC Biotechnol 19, 68. 10.1186/s12896-019-0558-y.

